# Jasmonate restrains the enhanced growth response of lateral roots induced systemically by a local water deficit

**DOI:** 10.64898/2026.10.05.756752

**Authors:** Xuelian Li, Virginia Protto, Christophe Maurel, Philippe Nacry

## Abstract

Water availability in drying soils is most often spatially heterogeneous, requiring plants to adjust their root system architecture to sustain water uptake. This acclimation response to water deficit (WD) relies on the integration of local and systemic signals. The later and their mode of action remain poorly understood. Using hydroponic split-root systems in maize and Arabidopsis, we show that, with respect to control well-watered plants, a local WD does not affect shoot biomass but suppresses lateral root (LR) growth in the water-deprived compartment while promoting LR elongation in the well-watered compartment. Spatiotemporal analyses in maize revealed an early and transient (<2h) transcriptomic response of LRs triggered by both local and systemic WD signals. This response involved activation of jasmonate (JA) biosynthesis and signaling components, including MYC2 like regulators and JAZ repressors. It occurred in both the water-deprived and well-watered root halves of split roots, with no evidence of long-distance transport of JA. Furthermore, exogenous methyl jasmonate locally inhibited LR elongation, whereas the Arabidopsis JA insensitive *jar1-1* mutant showed an enhanced systemic LR growth response to a local WD. Together, these findings demonstrate that JA acts as an early negative regulator that counterbalances the systemic enhancement of LR growth induced by a local WD, thereby fine-tuning root system architecture and water uptake during local soil drying.

**Significance Statement:** Plants growing in heterogeneous soils must coordinate local and systemic responses to optimize water acquisition, yet the underlying signaling mechanisms remain poorly understood. Using hydroponic split-root systems in maize and Arabidopsis, we show that local water deficit rapidly suppresses root growth in the stressed root half while stimulating lateral root elongation in the unstressed half. Spatiotemporal transcriptomic analyses reveal a rapid, transient activation of jasmonate biosynthesis and signaling. Genetic evidence further demonstrates that jasmonate restricts systemic lateral root elongation, acting as a negative regulator of developmental plasticity. These findings uncover a balanced interplay between positive and negative effectors and an unexpected role for jasmonate as an early integrator of systemic root responses to heterogeneous water availability, expanding its established functions beyond defense and canonical drought signaling.

## Introduction

Water availability is recognized as the most limiting environmental factor for plant growth and agricultural productivity (1–2). This constraint is intensified by climate change, which drives more frequent and severe drought events. The losses in plant productivity and survival due to drought are therefore projected to double by 2080 (1–2). A closer inspection of natural and agricultural drying soils reveals that water availability around plant roots is highly heterogeneous. At the macro scale, limited rainfalls cause surface soil layers to preferentially dry due to evaporation and intense root water uptake, generating vertical gradients in soil moisture (3–5). Horizontal heterogeneity also arises from plant competition and localized differences in soil water retention (6). At finer scales, heterogeneity in soil structure and composition leads to uneven water distribution, with features such as macropores generating localized zones of severe water deficit (WD) (see extensive reviews by 7,8). Such non-uniform soil moisture patterns may definitely challenge the ability of roots to meet the shoot water demand. Yet, root systems can show dramatic acclimation responses to spatially variable water availability by adjusting their growth and thereby their architecture as well as their water transport (hydraulic) properties (9, 10). Understanding how roots sense, integrate, and respond to heterogeneous water availability is of prime fundamental interest and essential for advancing agronomic research and sustainable agriculture.

Recent studies have revealed the molecular mechanisms enabling plant roots to perceive local moisture gradients and adjust their growth accordingly. Roots can redirect elongation or initiate new lateral roots (LRs) toward zones of higher water availability through processes known as hydrotropism (11) and hydro-patterning (12, 13), respectively. Conversely, LR formation is locally suppressed in water deprived regions, such as air-filled soil macropores, through a process named xerobranching (14). Together, these responses ensure that local root growth and branching are preferentially positioned towards zones of higher water availability.

In addition to local root responses, plants display systemic responses to heterogeneous water availability. In agriculture, these systemic effects are exploited through irrigation techniques that intentionally create spatial and temporal soil moisture heterogeneity known as partial root-zone drying (PRD). These techniques trigger whole-plant signaling responses that help regulate the plant water status and improve water-use efficiency (15–18). Several studies have shown that PRD affects stomatal opening, plant hydraulic conductivity, root growth, and nutrient uptake. However, the molecular mechanisms underlying these responses remain largely unresolved (19–23). Yet, plant adjustment to PRD primarily relies on the developmental plasticity of the root system. Rice, for instance, was shown to alter its root system architecture (RSA) to maintain efficient water uptake under PRD conditions (24).

Hormones are thought to play a central role in the local and systemic signaling processes that regulate RSA under varying environmental conditions. Several hormones have been detected in the xylem and phloem sap and likely contribute to the coordination of systemic stress responses. In *Arabidopsis* for instance, abscisic acid (ABA), a key regulator of plant acclimation to water deprivation (recently reviewed by 25) has been shown to coordinate overall root growth and hydraulic responses under osmotic stress (26). Furthermore, ABA functions as a key signal in the establishment of hydrotropism and xerobranching (11, 14). Jasmonates collectively refer to jasmonic acid (JA) and its derivatives, methyl jasmonate (MeJA), and jasmonoyl-isoleucine (JA-Ile). Jasmonates are central regulators of plant growth, development, and responses to biotic and abiotic stresses, with growing evidence linking JA signaling to drought tolerance and root developmental plasticity (27–28). MeJA priming enhances drought resistance in several plant species (29). Endogenous JA and JA-Ile levels rise rapidly and transiently following drought exposure in *Arabidopsis* and citrus, with JA likely acting upstream of ABA signaling (30). In rice, the stress responsive gene OsJAUP1, triggers a feed-forward loop activating JA biosynthesis and signaling genes, thereby enhancing WD tolerance and modulating auxin-dependent root development (31). JA signaling can also operate systemically, as jasmonates are transported over long distance through vascular tissues (32, 33).

The biological actions of JA are tightly linked to the regulated biosynthesis of its bioactive derivatives. In brief, JA biosynthesis and the formation of its bioactive derivatives is a key feature of this hormone biological actions. In brief, linolenic acid is first converted into 12-oxo-phytodienoic acid (12-oxo-PDA) via the sequential action of LOX, AOS, and AOC enzymes. 12-oxo-PDA then undergoes reduction and three cycles of β-oxidation to produce JA (34). The bioactive conjugate JA-Ile is then generated by JAR1 (35, 36). Under low JA-Ile levels, transcription factors regulating JA-responsive genes remain repressed through the action of JAZ transcriptional inhibitors (37). Abiotic stresses stimulate JA-Ile accumulation in the cytosol and its translocation into the nucleus. There, JA-Ile promotes the interaction between COI1 and JAZ proteins which leads to JAZ degradation and activation of JA-responsive gene transcription (34, 37).

Although numerous studies have examined local effects of water availability on root tip or LR formation, our understanding of whole-root system growth under heterogeneous water availability remains limited. In particular, the functional integration of local sensing of WD with systemic signaling is still poorly characterized. In this study, we established a simplified split-root system to analyze root responses to a localized WD (osmotic stress) and explored the early signaling events underlying both local and systemic developmental regulation. Our results show that, with respect to plants grown under well-watered (WW) conditions, a local WD represses root development in the water-deprived root half while stimulating growth systemically in the water-sufficient root half. We further demonstrate that a local WD triggers a transient increase in jasmonate biosynthesis in the whole root system. We propose that jasmonate synthesized in the water-sufficient root half partially counteracts the enhanced growth of LRs in this half, thereby fine tuning the systemic root growth response to a local WD.

## Results

### A local polyethylene glycol (PEG)-induced WD systemically promotes LR development in both maize and Arabidopsis

In order to explore the local and systemic developmental responses of roots to a local WD, we developed a hydroponic split-root system where one half of the root system of maize or Arabidopsis plants can be subjected to a PEG-induced WD whereas the other half is bathing in a standard solution, yielding WW conditions (Fig. 1a).

**Figure 1.**
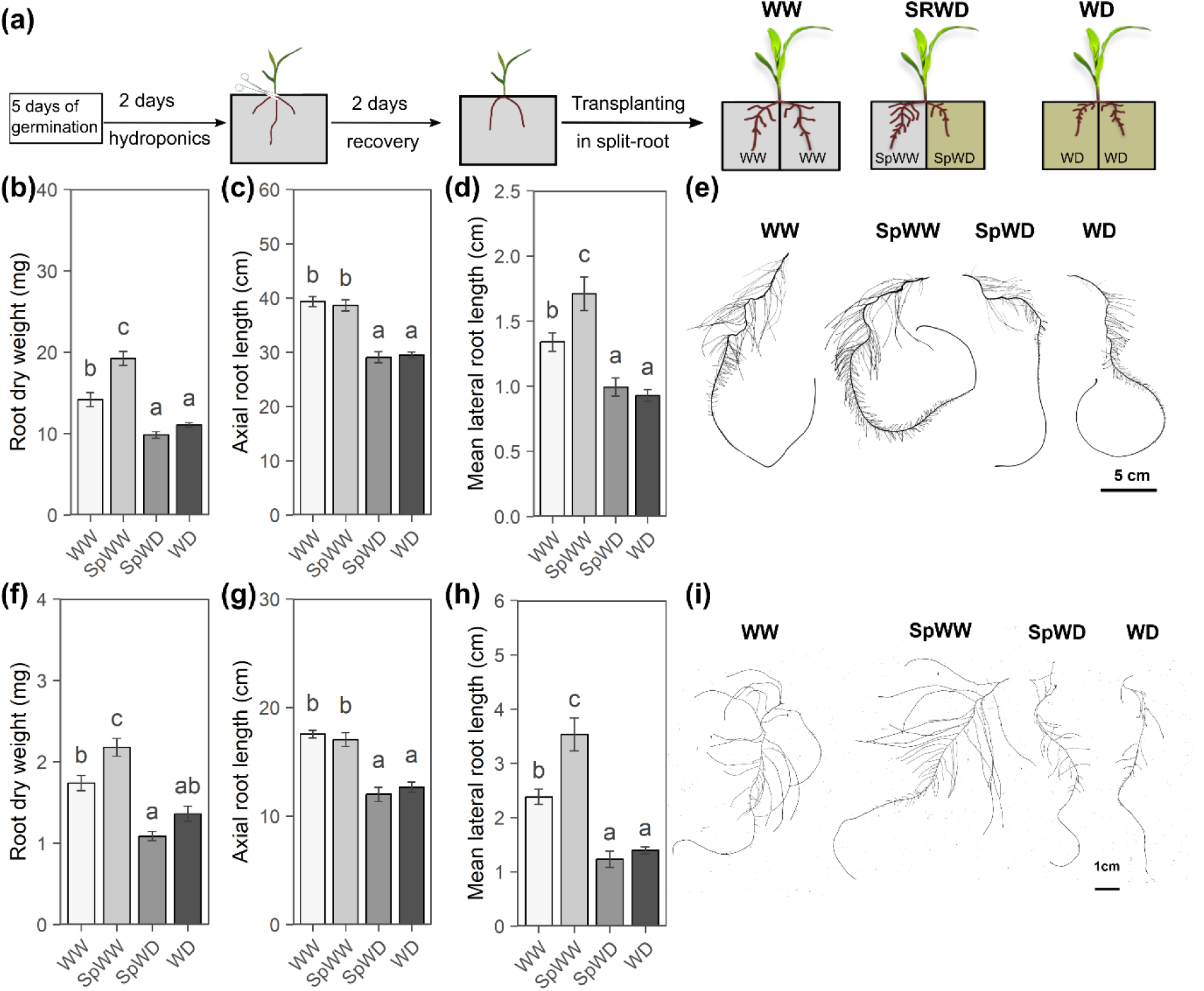
Growth responses of maize and *Arabidopsis* seedlings to a PEG-induced local WD. (**a**) Experimental design of a split-root hydroponic system to explore in maize seedlings the effect of a local WD on root growth and gene expression. WW: plants with two root halves grown in a standard hydroponic solution; WD: plants with two root halves grown in a standard hydroponic solution supplemented with 150 g L^-1^ PEG; SRWD: plants grown in the split root system with one half of the roots in the standard solution (SpWW) and the other half in the standard hydroponic solution containing 150 g L^-1^ PEG (SpWD). A similar set up was used for Arabidopsis. Root dry weight (**b**), axial root length (**c**) and LR growth (**d**) of maize seedlings grown for 3 days under standard conditions (WW), a homogeneous (WD) or a local (SpWD) WD. (**e**) Representative images of maize root development in response to these growth conditions. Root dry weight (**f**), axial root length (**g**) and LR growth (**h**) of *Arabidopsis* after 4 days under standard conditions (WW), or a homogeneous (WD) or local (SpWD) WD. (**i**) Representative images of Arabidopsis root development in response to these growth conditions. The bar plots show cumulated data from 3 independent experiments (n=16-18 plants). All data represent mean ± SE. For each root type, different letters indicate statistically different values (P < 0.05, one-way ANOVA followed by FDR test).

In maize, a homogeneous WD (−0.28 MPa) reduced after 3 to 5 days both shoot and root growth (dry weight) by 22 to 32 % (Fig 1a and b; Fig.S1a ; Fig. S2a and b). By contrast, plants subjected for the same time to a local WD (Split WD: SpWD) had the same shoot dry weight as WW plants (Fig.S1a and Fig. S2a). Yet, these plants showed a markedly modified root growth. At 3 days, the SpWD root halves showed a tendency to growth inhibition as compared to WD root halves, with a 11.7% lower dry weight (Fig. 1b). Conversely, the SpWW root halves showed higher growth than WW root halves, with a 35.2% higher dry weight (Fig. 1b). These responses were amplified after 5 days with a significant reduction of growth (−35% dry weight) in SpWD *vs.* WD root halves and a higher enhancement of growth (+44% dry weight) in SpWW *vs.* WW root halves (Fig. S2b). Analysis of RSA revealed that 3 days of a homogeneous or heterogeneous WD had same local effects, and similarly reduced axial (seminal) root length by 25% (Fig. 1c). LR development, as analyzed through total length, average length, number, or density was also inhibited locally, in a same manner by a homogeneous or heterogeneous WD (Fig 1d, Fig S1 c-d). On the contrary, SpWW root halves had longer LRs (Fig. 1d; Fig. S1b). Five days of a split WD further amplified the differential growth of SpWW *vs.* WW root halves (Fig. S2d). Furthermore, LR emergence was enhanced in SpWW compared to WW, while it was more strongly inhibited in SpWD compared to WD root halves (Fig. S2e), resulting in a characteristic imbalance in RSA (Fig. 1e). To assess whether these responses are conserved across species, we subjected *Arabidopsis* plants to a four-day split WD in hydroponics. Similar responses to those of maize were observed for shoot and root dry weight and RSA (Fig. 1 f-I; Fig. S1 e-h). Overall, split WD induced a marked increase in LR growth in SpWW root halves compared with plants grown under homogeneous WW conditions indicating that the systemic stimulation of LR elongation is a conserved response.

### Transcriptomic analyses reveal dynamic and organ specific responses to a local WD

We designed targeted transcriptomic experiments in maize to investigate the molecular mechanisms underlying the systemic responses of roots to a local WD. For this, we sampled two root fractions: the unbranched zone of axial (seminal) roots from the tip to the first visible LR (Tip) and the LRs, with 9 time points ranging from 0.5h to 96h after PEG treatment (Fig S3 a). LR and Tip samples from plants grown under WW, homogeneous WD or split WD were collected in four replicates, generating a set of 296 transcriptomes. Global analysis of all samples revealed expression of 26148 genes with average read counts over 10. Principal Component Analysis (PCA) of all expressed genes indicated that Dimension 1 (16.7% of variance) separated samples according to WD duration, while Dimension 2 (13.4%) distinguished LR and Tip transcriptomes (Fig. S3a–c) which, therefore, were analyzed independently (Fig. S3 f-k). In both cases, WD and SpWD samples grouped together as did WW and SpWW. In addition, WD duration massively altered gene expression. When WD treatments were categorized as short (0.5h-4h), intermediate (8h-24h), and long (48-96h) (Fig. S3f and i), LR transcriptomes displayed a clearer separation on axis 1 according to treatment duration (Fig. S3f) than Tip transcriptomes (Fig. S3i). These analyses are consistent with the differential developmental responses of SpWD and SpWW root halves with respect to their WD or WW counterparts, which were more pronounced in LRs than in Tip samples.

We then used the WD *vs.* WW comparison as a positive control for exploring further the experiment robustness. We identified 1651 and 998 differentially expressed genes (DEGs) in response to WD in LRs and Tips, respectively, with FDR ≤ 0.01 and |log2FoldChange| ≥1 (Fig 2a-b). Among them, 47.2% were specific to LRs and 13.4% to Tips while the remaining DEG (39.4%) were common to both types of samples. To identify genes putatively involved in the systemic control of root response to a split WD treatment, we then searched for DEGs in SpWW *vs*. WW and SpWD *vs.* WD comparisons. When considering all time points together, 65 and 57 DEGs were identified between SpWW and WW and 78 and 7 between SpWD and WD, in LRs and Tips, respectively (Fig. 2a-b).

**Figure 2.**
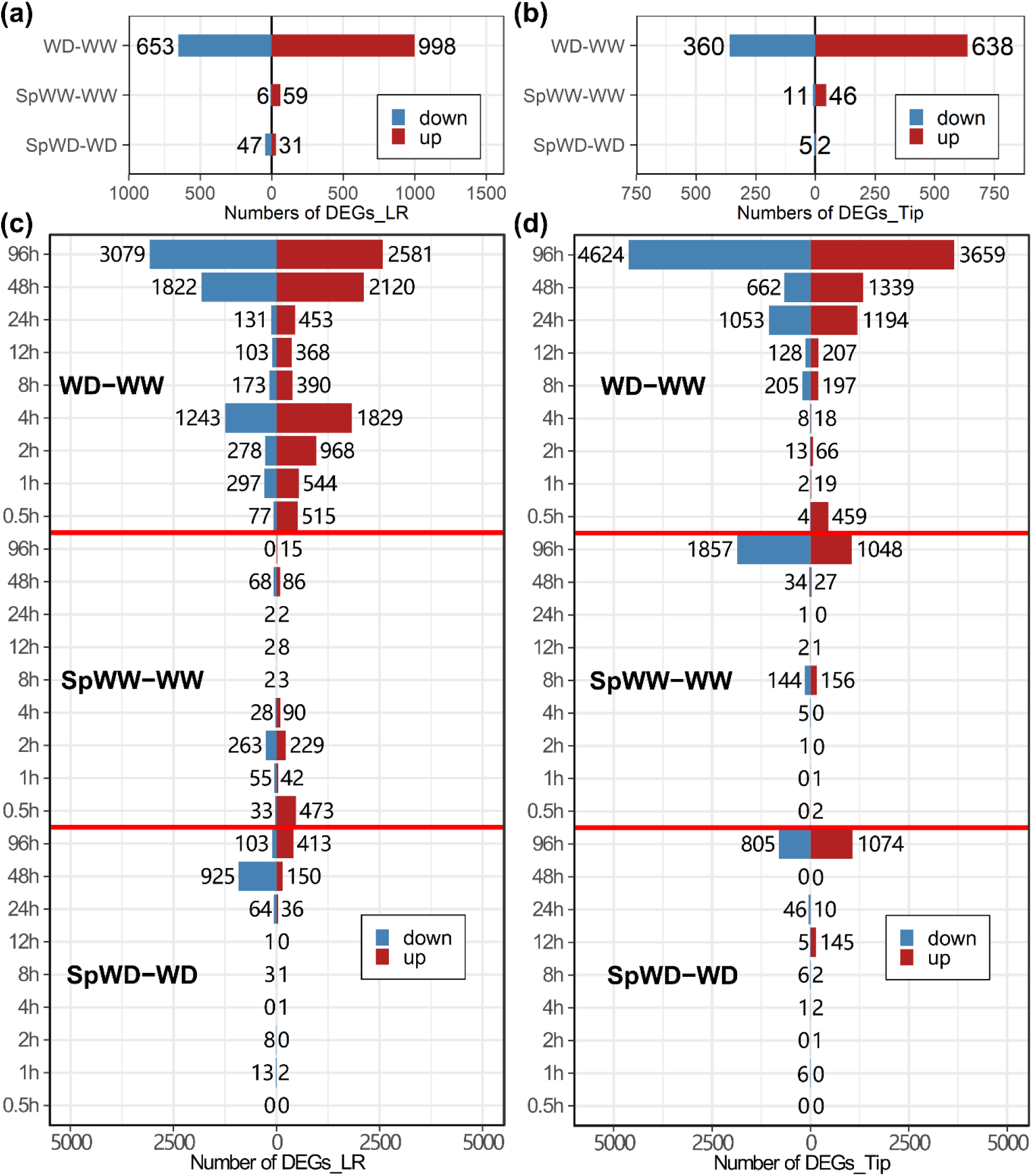
Transcriptional responses of maize roots to PEG-induced local WD. (**a**, **b**) Number of DEGs in LR (a) and Tip (b) samples, respectively, of maize seedlings, after a homogeneous (WW,WD) or heterogeneous (SpWW, SpWD) treatment. (**c**, **d**) Number of DEGs in LR (c) and Tip (d) samples at indicated time points. The figure compares roots of plants subjected to the same treatments as above. DEGs were identified using DESeq2 with P <0.01 and |Log2 fold change|>=1 thresholds. The figure shows gene expression data from 4 independent biological (plant culture) experiments.

Because these numbers of DEGs are low, we then considered individual time points to possibly reveal dynamic and transient molecular responses. For reference, a homogeneous WD induced a massive response in both the early (0.5-4h) and late (48-96h) stages in LRs (Fig. 2c), whereas Tip samples only exhibited a late response (8-96h) (Fig. 2d). Regarding systemic responses, a split WD deregulated ∼600 genes within 0.5-4h in untreated LR samples (SpWW *vs.* WW) whereas differential expression was essentially observed after 48h in PEG-treated LRs (SpWD *vs.* WD) but with a higher number of DEGs (∼ 1300 genes) (Fig. 2c). By comparison to LRs, a high number of DEGs (∼2900) was observed in Tip samples only at later time points in the same conditions (SpWW-WW: 96h; SpWD-WD: 96h) (Fig. 2d). In summary, our findings indicate significantly different profiles in the responses of LRs and Tip to WD, with LRs responding quickly and massively to a homogeneous WD, while the Tips part mainly responds after a 4-8 h delay. The early and transient molecular responses in the SpWW root halves reveal a systemic response that we attribute to a long-distance signaling process, from SpWD towards SpWW root halves, these responses being faster in LRs than Tips tissues. Finally, the detection of numerous DEGs between SpWD and WD after 48-96h in both LRs and Tip suggests another longer-term acclimation response, specific to split WD. Overall, these results point to time-and root type-dependent systemic events that may collectively shape the integrated response of root halves to a split WD.

### A unique set of transcription factor genes is transiently induced in LRs by a WD, both locally and systemically

Based on the results above, we reanalyzed the LR transcriptomes integrating time as a cofactor. Eight groups were formed by k-means clustering (Fig. 3a). Groups 1 to 3 contain genes mainly expressed in WW and SpWW root halves (Fig S4a) and with ontologies related to cytoskeleton structure, cell wall organization, mitotic cell cycle, H2O_2_/ROS metabolism, antioxidant activity, and water transport (Fig. S4b-d; Table S1-2). Groups 5 to 8 contain genes that are upregulated in both WD and SpWD halves (Fig. 3a; Fig. S4a). The putative functions of these genes are related to defense response, fatty acid elongation and biosynthesis of secondary metabolites (Fig. S4f-i; Table S1-2). Group 4 is unique in that the expression levels of its 210 DEGs increased dramatically and transiently in WD, SpWD, and SpWW LR samples shortly (< 30 min) after transfer to split-root conditions, whereas this response was delayed and greatly reduced in WW (Fig. 3a; Fig. S4a). This unique expression profile identifies genes responding similarly to a local WD stimulus or to a systemic stimulus transferred from SpWD to SpWW root halves. Remarkably, 47 of the 65 DEGs detected in the global analysis between SpWW *vs.* WW samples (Fig 2a) are present in this group (Fig.3b). Gene Ontology (GO) analysis identified “JA mediated signaling” as the most significantly enriched GO term among Group 4 genes (Fig. S4e; Table S2).

**Figure 3.**
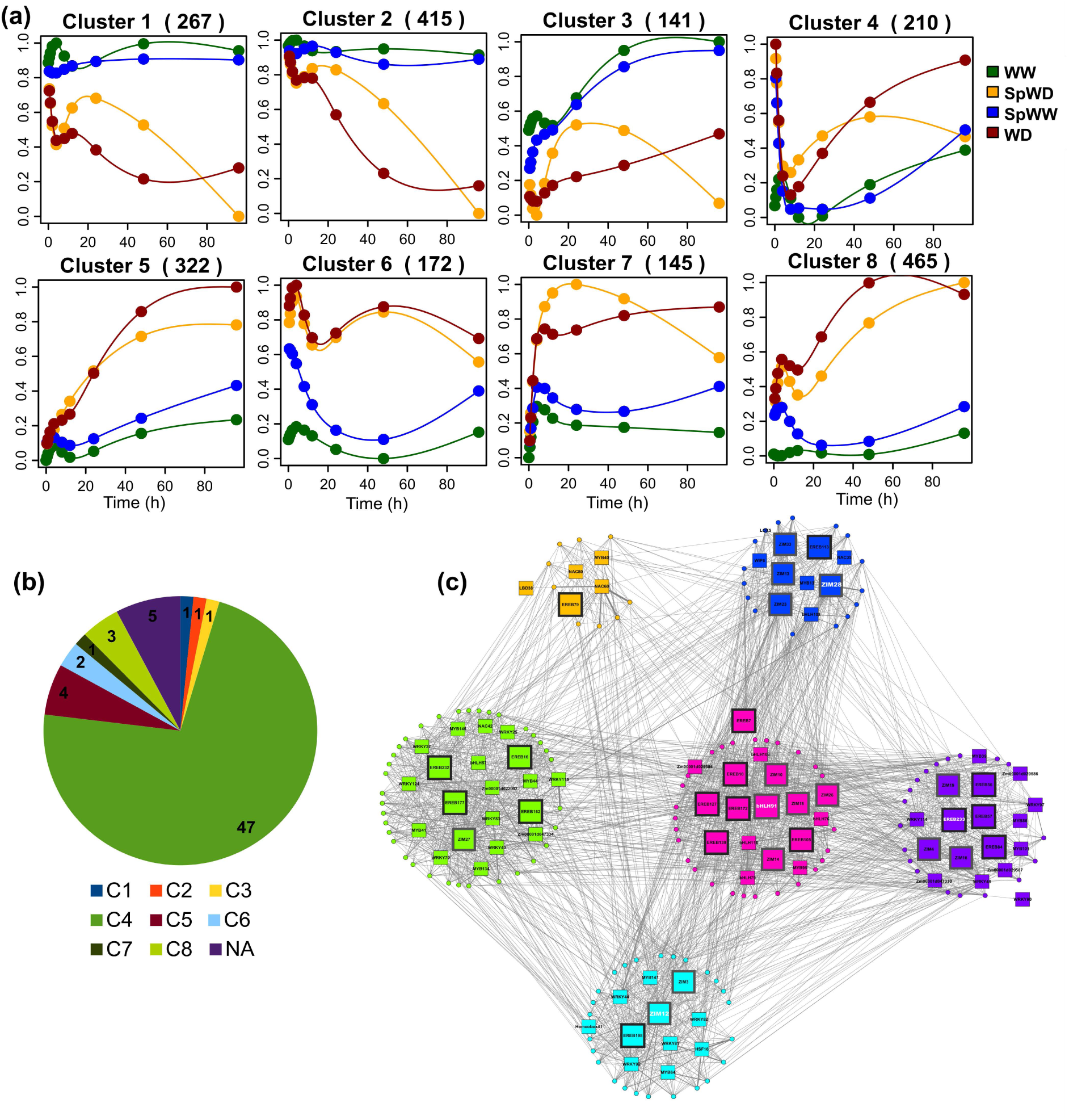
Identification of a gene cluster rapidly and transiently induced by a local WD in LRs, both locally and systemically. (**a**) k-means clustering analysis, focusing on DEGs identified in LR in response to a heterogeneous WD, identifies eight major DEG groups. The number of genes in each group is mentioned in brackets. The lines represent the scaled gene expression for WW (green), SpWW (blue), SpWD (orange) and WD (brown) samples. (**b**) Repartition within the different groups of the 65 DEGs identified in the SpWW *vs.* WW comparison (Fig.e 2a); the number of DEGs in each group is mentioned inside the partition of the pie chart. (**c**) Gene regulatory network built with the genes of Group 4. Square symbols represent TF regulators, and circles represent target genes. The larger and bolder squares identify jasmonic acid-and ethylene-responsive TFs. *ZmbHLH91, ZmEREB233, ZmZIM12* and *ZmZIM28* were highlighted in white. Nodes in a same color identify subgroups with stronger expression correlation.

Subsequently, we constructed a gene regulatory network (GRN) based on Group 4 by incorporating regulators from the “planttfdb” database (http://planttfdb.gao-lab.org/index.php?sp=Zma) and the transcription factors filtered with “BioMart” under the EnsemblPlants database “Ensembl Plants Genes 49 - Zea mays genes (B73 RefGen_v4)” (Table S3) (Fig. 3c). The 210 DEGs of Group 4 fall into 78 regulators and 132 target genes. These regulators include 69 transcription factors (TFs) belonging in particular to the AP2-EREBP/ERF, ZIM, WRKY and MYB classes (Table S4). Most of them have been documented to respond directly or indirectly to JA (38). Interestingly, the JA signaling master regulator *ZmbHLH91* (Zm00001d047017), an ortholog of the *AtMYC2* (39), was identified as one of the most interconnected TF gene (Fig. 3c). In addition, many AP2-EREBP/ERF TFs were also functionally annotated as ethylene (ET) responsive TFs (Table. S1). These findings support our observation that homogeneous and local WD treatments result in LR-specific short-term transcriptional regulations, that occur both locally and systemically. They indicate that JA and/or ET could be key regulatory hormones in these processes.

### Split WD systemically induces a JA dependent signaling pathway in maize LRs

We then focused on six putative JA-and ET-responsive TFs (*ZmEREB79*, *ZmEREB233*, *ZmbHLH91*, *ZmZIM12*, *ZmZIM27* and *ZmZIM28*), selected as representative markers of Group 4. Profiles extracted from transcriptomic data showed that their expression was rapidly induced in response to a local WD, both locally and systemically, with a first peak occurring 0.5-1 h after treatment (Fig. S5 a-f). The rapid local induction by a WD was confirmed by RT-qPCR in an independent set of experiments (Fig. 4a). We also observed that the six TF genes were induced in LRs after 1h of exposure to 50 µM MeJA (Fig. 4a). Yet, only *ZmEREB79* and *ZmEREB233* were induced after 1h by a root treatment with the ET precursor 1-amino-cyclopropane-1-carboxylic acid (ACC) (Fig 4a and Fig. S6 f). Given the similar expression profiles observed among the 6 TFs, only *ZmZIM28* (Fig. 4b, c) and *ZmZIM12* (Fig. S5g, h) were selected for further analyses. We observed that their expression was also transiently induced in shoots following exposure of roots to a local or homogeneous WD, although to a much lower extent than in LRs. More generally, the MeJA-induced expression of *ZIM12* and *ZIM28* was dose dependent and remained stable after 4h (Fig. S6 a-b) whereas little or no induction was observed in response to a wide range of ACC concentrations (Fig. S6 d-e). Interestingly, when combined with WD, exogenous MeJA did not over induce the *ZIM28* and *ZIM12* but the highest concentrations (10 and 50 μM) maintained their high expression over time (Fig. S6 a-b). These results indicate that the initial molecular response to WD was more likely influenced by JA-dependent than ET-dependent transient signaling processes.

**Figure 4.**
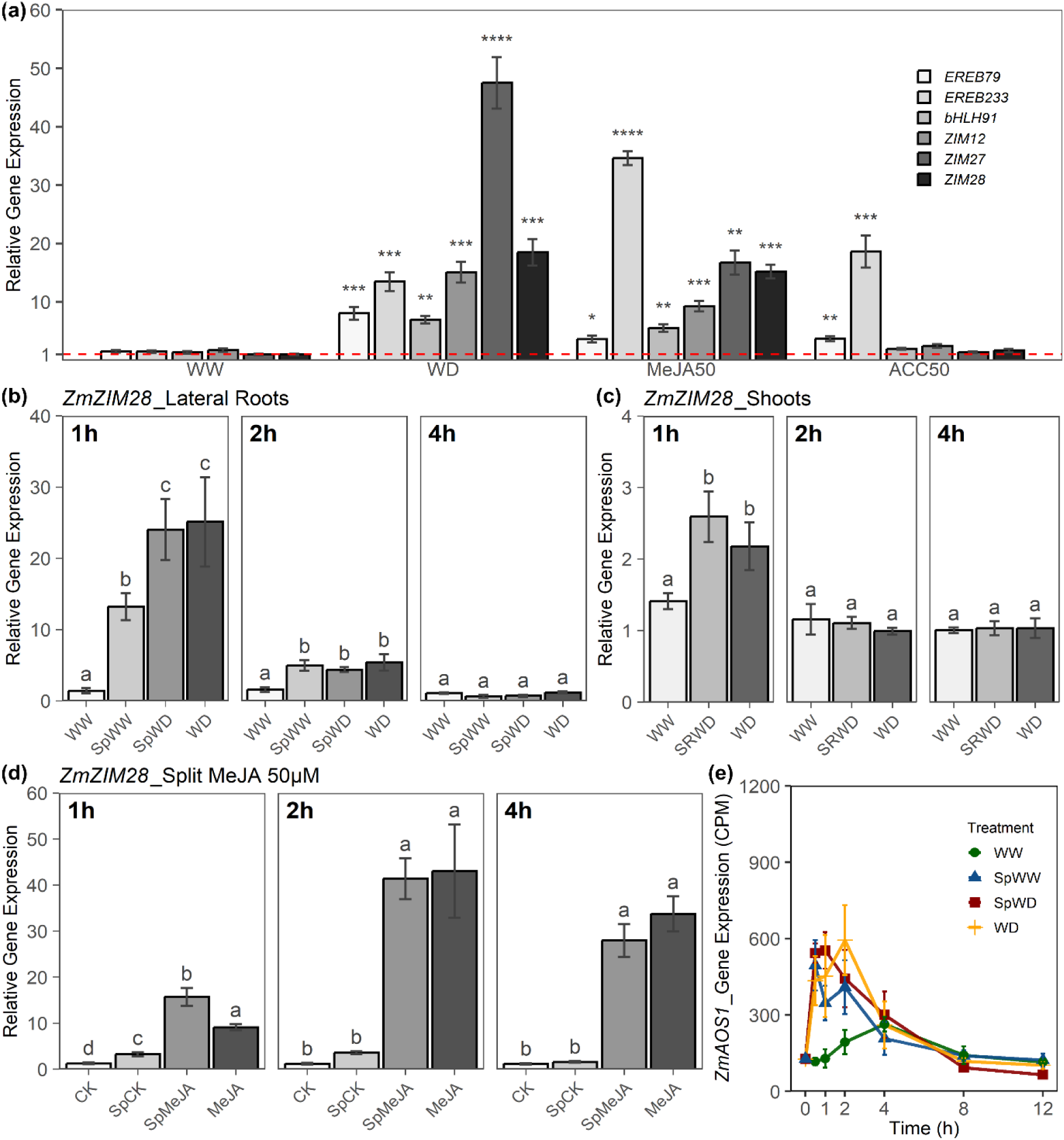
Expression pattern of JA responsive genes in maize LRs. (**a**) Expression pattern in LRs of several *ZmZIMs* and *ZmEREBs* selected from Group 4 after 1h of control (WW), PEG (150 g/L) induced WD, 1h methyl-jasmonate (50 µM -MeJA50) or 1h 1-aminocyclopropane1-carboxylic acid (50 µM - ACC50) treatments and compared to control culture condition before treatment. Gene expression was monitored by RT-qPCR (**b-c**) *ZmZIM28* expression in LRs and shoot in response to PEG induced local WD; (**d**) *ZmZIM28* expression in LRs after a short term homogeneous or local MeJA (50uM) application; (**e**) Expression pattern of JA biosynthesis gene *ZmAOS1* in LRs in response to a local WD. Experimental conditions and figure conventions for local WD are as above. For split-root MeJA treatment, one half of the roots was grown in a standard solution (SpCK) and the other half in a standard hydroponic solution supplemented by 50 µM MeJA (SpMeJA). CK and MeJa correspond to plants with a standard solution or a MeJa-supplemented solution, respectively, applied to their two root halves. All data represent mean ± SE of 4 biological replicates with 2 technical repeats; Asterisks indicate differences (Student’s t test; *P < 0.05; **P < 0.01; ***P < 0.001; ****P < 0.0001) with WW in different genes, respectively; For each time point after treatment, different letters indicate statistically different values (P < 0.05, one-way ANOVA followed by FDR test).

To test the hypothesis of a long-distance transport of JA, acting from root-to-shoot-to-root, 50µM exogenous MeJA was applied to one side of roots under split-root hydroponic (SRMeJA) conditions. The expression of *ZmZIM28* and *ZmZIM12* was strongly and continuously induced in LRs of SpMeJA root halves (Fig. 4d; Fig. S5i). The induction was similar to that observed in LRs from a homogeneously treated root system (MeJA). By contrast, no induction of expression was observed in the LRs of complementary control split-root (SpCK) root halves (Fig. 4d; Fig. S5i). Similarly, no significant overexpression was observed when MeJA was sprayed onto the leaves (data not shown). These results suggest that JA is poorly transported at long distance and that the systemic signaling observed in response to split WD may rather rely on local JA biosynthesis. When exploring transcriptomic data, we observed indeed that most of the genes involved in JA biosynthesis were rapidly and transiently induced by a homogeneous or local WD. Independent Q-RT-PCR experiments confirmed that these genes were induced systemically and massively in the control root half of WD-treated split roots (Fig. 4e; Fig. S7). These data further support the hypothesis that JA was not significantly translocated from the root half under WD to the unstressed half, but was rather biosynthesized locally in both root halves.

### Split WD induces a systemic JA-dependent signaling pathway limiting LR elongation

In order to explore the direct impact of JA on RSA of maize and Arabidopsis, we applied a wide range (0.1-50 µM) of exogenous MeJA, a volatile methyl ester of JA that can be converted back to JA in plant tissues by esterases, to the roots of seedlings of both species. Increasing concentrations of MeJA inhibited both axial and LR elongation in a dose-dependent manner (Fig. S8). These results challenge the idea that systemic induction of JA biosynthesis in the SpWW half of split roots under WD could account for the typical enhancement of LR elongation. To better explore the role of JA signaling in root adaptive responses to a local WD, we used a molecular and genetic approach in *Arabidopsis*. Amino acid alignments identified *AtJAZ2* and *AtJAZ1* as the best-fitting Arabidopsis orthologs of *ZmZIM28*. Expression of *AtJAZ2* and *AtJAZ1* was strongly and transiently induced by a short-term homogeneous application of MeJA (50 µM) (Fig.5a; Fig. S9a). Their expression was also transiently and massively induced under a homogeneous (Fig.5b; Fig. S9b) or local WD, both locally and systemically (Fig.5b; Fig. S9b) making *AtJAZ2* and *AtJAZ1* good reporters for further analyses. In the roots of the JA signaling-deficient mutant *jar1-1* (40), expression of *AtJAZ2* and *AtJAZ1* was significantly lower than in Col-0 (Fig.5c; Fig. S9c). Notably, neither a homogeneous nor a local WD induced their expression regardless of the treatment duration (Fig.5c; Fig. S9c).

Based on these results, we analyzed the RSA responses of *jar1-1* to these treatments (Fig.6a). Col-0 and *jar1-1* both exhibited a similar root and shoot dry weight under WW and homogeneous WD conditions (Fig.6b; Fig. S10a). In split-root experiments, both genotypes showed similar DW of their shoots (Fig. S10a) and SpWD root halves (Fig.6b). By contrast, the DW of SpWW root halves was higher in *jar1-1* than in Col-0 (Fig.6b). In these root halves, *jar1-1* showed a 31% increase in total root length (Fig. S10b) compared to Col-0 due to an overstimulation of both axial (1^st^ order LRs) and LR (2^nd^ order LRs) length (Fig. 6 c-d), but not to the production of supernumerary LRs (Fig. S10 c-d). The overall data show that JAR1 mediates a partial repression of the enhanced growth of SpWW root halves whereas no significant role can be revealed in root growth inhibition induced by a local WD treatment in SpWD or WD root halves.

**Figure 5.**
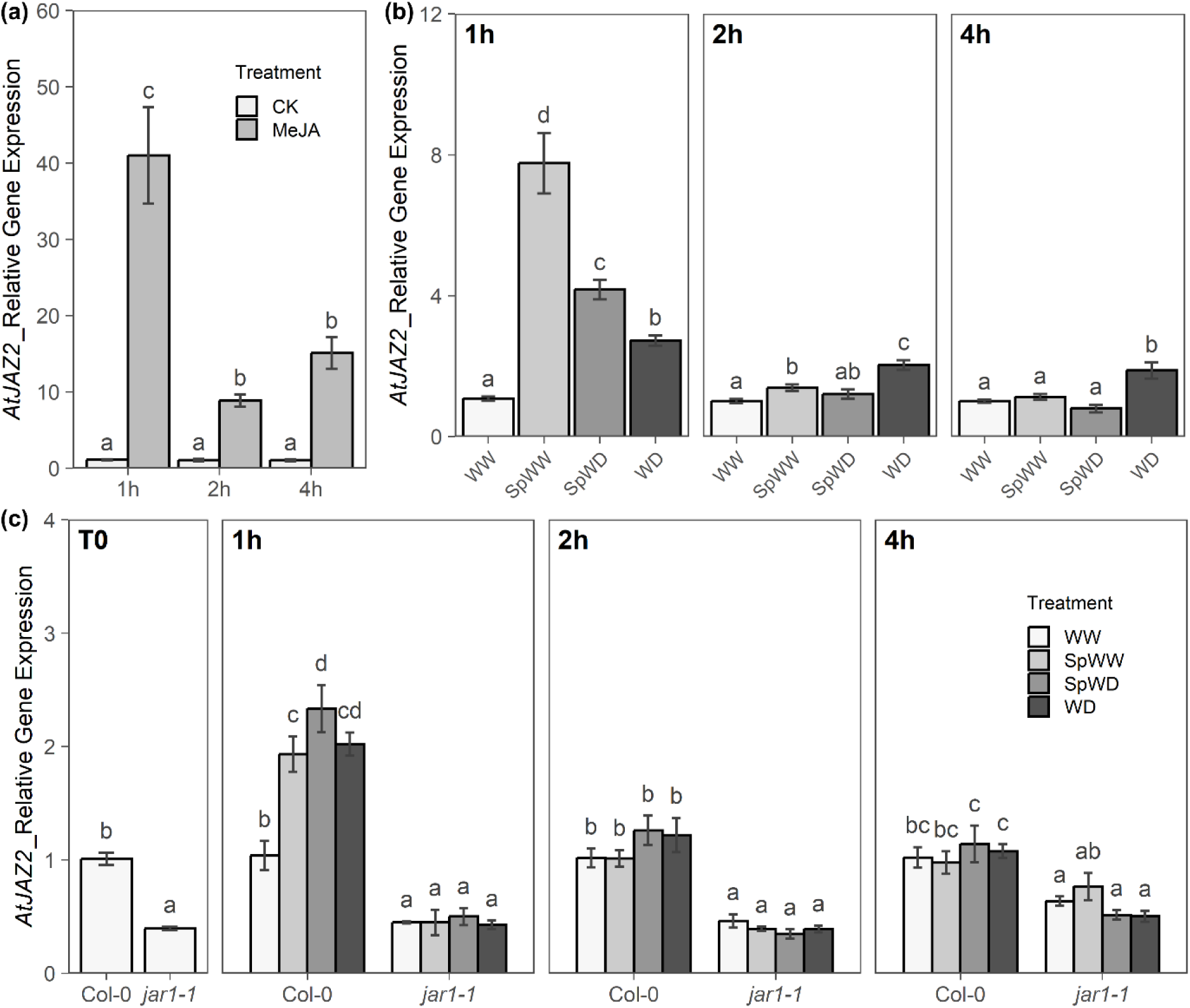
Expression pattern of JA-responsive gene *AtJAZ2* in *Arabidopsi*s. (**a**) Kinetic expression profile in Col-0 roots of *AtJAZ2* in response to a 50 µM MeJA treatment; (**b**) Kinetic expression profile in Col-0 roots of *AtJAZ2* in response to a local WD; (**c**) Kinetic expression profile of *AtJAZ2* in Col-0 and *jar1-1* roots in response to a local WD. Gene expression was monitored in Arabidopsis roots by RT-qPCR. Experimental conditions and figure conventions for local WD are as above. All data represent mean ± SE of 3-4 biological replicates with 2 technical replicates. For each time point after treatment, different letters indicate statistically different values (P < 0.05, one-way ANOVA followed by FDR test).

**Figure 6.**
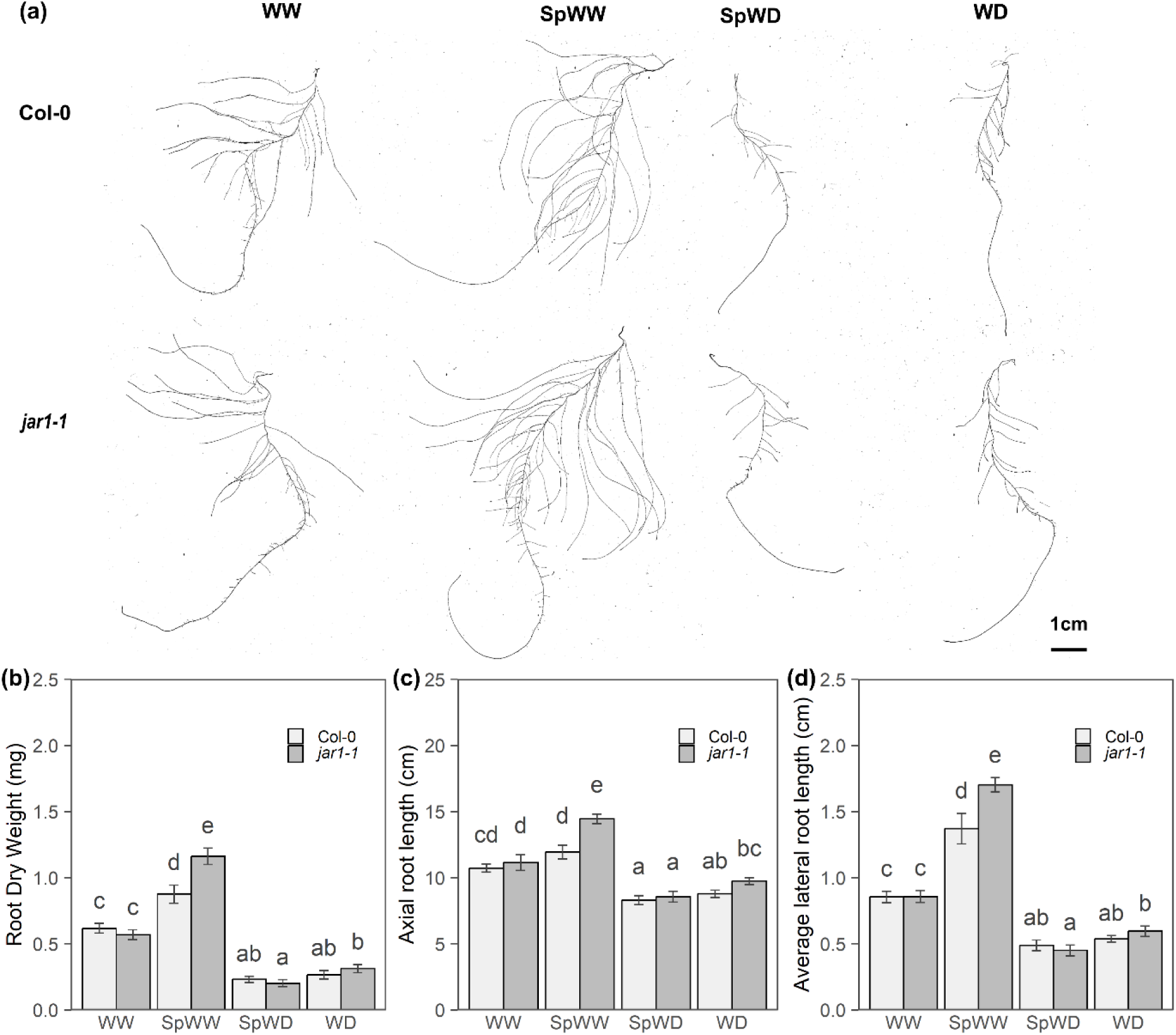
Growth responses of *Arabidopsis* Col-0 and *jar1-1* seedlings to a local WD. (**a**) Representative images of root development in response to homogeneous or local WD for 4 days; (**b**) Root dry weight and (**c-d**) Axial root length and average lateral root length after 4 days of growth. Experimental conditions and figure conventions for local WD are as above. The bar plots show cumulated data from 2 independent experiments (n=12-20). All data represent mean ± SE. For each root trait, different letters indicate statistically different values (P < 0.05, one-way ANOVA followed by FDR test).

## Discussion

In the present work, we developed a split root system using PEG to modify water availability at macroscopic level. With respect to a soil system, our set up allows to synchronously modify the environmental conditions while providing an easy access to the roots. Using hydroponic cultures, this system enabled us to observe a marked enhancement of LR elongation in the WW root half without any change in LR density (Fig.1 and Fig.S1), in both maize and Arabidopsis (Fig.1). This response is clearly distinct from localized responses to water availability namely hydropatterning (11) and xerobranching (14) that involve the initiation and early development of LR primordia. By contrast, it is similar to those reported in many different species in soils under partial root zone drying (reviewed by 10) and likely reflects a conserved mechanism in plants.

The split-root phenotypic responses were used to design a spatiotemporal transcriptomic analysis of local and systemic responses to a local WD. In good agreement with the marked growth responses of LRs, their transcriptional response occurred earlier (< 4h) and involved a larger number of DEGs than in axial roots. Based on kinetic gene expression profiles, we were able to identify homogeneous gene groups and showed that, in most of these, the WW and SpWW profiles on the one hand, and the WD and SpWD profiles on the other hand were largely related (Fig. 3a). Thus, root responses to water availability are essentially local. In agreement with this, the number of DEGs in LRs between the WD and SpWD conditions or the WW and SpWW conditions were dramatically lower (65 and 78, respectively) than in WW *vs.* WD comparisons (1651) (Fig. 2a). Nevertheless, kinetic modeling of gene expression using spline regression approach enabled us to identify a single group (Group 4) of 210 genes with a differential expression profile between SpWW and WW conditions. Interestingly, these genes showed a marked induction in SpWW root halves in < 30 min which then vanished (Fig. 3a). Although this early induction was much reduced in WW root halves, a puzzling observation was that it was also prominent upon local WD treatment, in LRs of WD and SpWD root halves.

Further analysis of the Group 4 genes revealed molecular functions associated with JA signaling (Fig. 3b). In maize, the ZIM family of JA-related TF consists of 49 members including 38 JAZ transcriptional repressors (41). We independently checked using qRT-PCR, that several of them including ZmZIM28 responded to local or homogeneous WD treatments, and showed rapid and marked induction in LRs of SpWW, SpWD and WD root halves. Interestingly, another JA master regulator, ZmbHLH91 (Zm00001d047017), an ortholog of the AtMYC2, that appeared as one of the most interconnected TF in our gene network (Fig 3c), showed the same transient induction expression pattern (Fig. S6d). This TF is known to mediate various crucial biological processes such as plant growth and stress responses (42). These results could be extended to Arabidopsis. The two close homologues of ZmZIM28, AtJAZ1 and AtJAZ2, exhibited similar induction by exogenous JA in LRs of SpWW, SpWD and WD root halves. Moreover, we showed these inductions to be under the control on JAR1 that conjugates JA into its bioactive form JA-Ile.

Several root sourced phytohormones, including ABA, ethylene, auxin and cytokinin have already been reported and extensively studied as mediators of root-to-shoot signaling, particularly in response to WD (43). Long-term effects of JA and other jasmonates have also been identified in roots of plants grown in homogeneously drying soil (44) or in response to PRD in cotton (45) and soybean (46). An early JA response, as identified here in both maize (Fig.3) and Arabidopsis (Fig.5), may have been overlooked in these previous studies, as they were conducted over longer time intervals. In particular, analyses in soil are usually performed after several days of treatment so that the PRD is well established. By contrast, and in line with our results, Luo et al. (45) used PEG to mimic a PRD in cotton and observed a maximal accumulation of JA at 1 hour after local application of PEG.

Although JA plays a key role in rapid long-distance signaling (within seconds) in response to defoliating herbivores, for example (32), this phytohormone has rarely been proposed as a major regulator of early signaling in the context of heterogeneous WD. The JA-dependent transcriptional responses observed in both maize (Fig. 3-4) and *Arabidopsis* (Fig.5) within minutes after treatment application, in SpWW but also SpWD and WD root halves, suggested to us that the hormone may be transported systemically via shoots to the hydrated root half (SpWW). Such systemic transport mechanism has been proposed by Luo et al. (2019) in cotton (45). However, experiments with local applications of JA to root or shoots parts did not support this hypothesis in maize. Transient induction of JA biosynthesis genes such as *ZmLOX4*, *ZmLOX5*, *ZmLOX8*, *ZmAOS1*, and *ZmAOS3* in the SpWW part further suggests that JA is synthesized locally (Fig. 4e; Fig. S7a-f). This hypothesis is consistent with the results reported by Puértolas et al., (2025) in soybean under PRD (46), who observed a higher accumulation and export of JA in the xylem sap of the wet part of the root system than in the xylem sap of the dry part. Collectively, these findings support local JA biosynthesis and acropetal transport. However, it remains to be determined whether cotton employs a comparable mechanism, exhibits a species-specific response, or whether prolonged local water deficit activates a feedback-driven basipetal transport of JA.

In the present work, we used treatments with exogenous JA to show that this hormone exerts a strong, dose-dependent inhibition on primary and LR growth and development, in both maize and Arabidopsis (Fig. S8), consistent with earlier literature (27, 40). Actually, effects of JA in plant responses to WD can be multiple. In particular, JA can attenuate ABA effects through antagonistic interactions, during stomatal closure (47) or root growth inhibition (48). Muzaffar et al. (2024) recently identified a positive regulator of JA biosynthesis in rice, JAUP1, which promotes root development and tolerance to several abiotic stresses by triggering ABA responses (31). When studying the response of cotton to PRD, Luo et al (2019) observed an accumulation of both JA and ABA in the dry part of the root system (45). Similarly, our data show, in WD and SpWD, a strong induction of genes involved in ABA biosynthesis and signaling (mainly present in Group 8) concomitant with the induction of JA biosynthesis and response genes (starting from 24 hours in WD and SpWD in Group 4). Interestingly, the latter induction is less pronounced in SpWD than in WD: this could explain the tendency toward an over-repression of root development in SpWD compared to WD, in both maize and *Arabidopsis* (Fig. 1 and Fig. S2).

While the accumulation of JA is consistent in the WD half of the maize root system, it may appear paradoxical in the WW half, given its inhibitory effects. Indeed, JA generally inhibits root growth and development (27, 40; Fig. S8a–h), whereas PRD (10, 49) and our split-root treatment promote LR elongation in wet roots (Fig. 1).Thus, we propose that the systemic activation of JA signaling in the wet root part acts as a negative feedback mechanism aimed at restricting development of the root system. An overdevelopment of LRs, combined with enhanced water uptake, as observed in cotton (45) and riparian Melaleuca species (21), could accelerate water consumption and possibly deplete the water resources of the wet soils area, thereby threatening plant survival. More generally, the fine interplay between positive and negative regulators is a common strategy that confers robustness to plant physiological processes. Such balancing mechanisms are well documented in plant immune responses where they play a crucial role in preventing excessive defense responses that could otherwise lead to growth inhibition, cell death, or even plant death (50).

In conclusion, the present work emphasizes the relevance of split-root hydroponic systems to explore the molecular mechanisms of local and systemic signaling in response to localized WD. We show that the fine and robust control of critical root growth responses implies a balanced interplay between positive and negative effectors. We further show that a rapid and systemic transcriptional control of JA-dependent signaling mechanisms limits root growth in the wet roots. This counterbalancing effect emphasizes the existence of unexpected mechanisms controlling LR elongation in roots under partial WD. Yet, the mechanism counteracted by JA that primarily enhances LR growth in SpWW roots remains to be identified. Since JA is not transported systemically, the long-distance signal(s) that trigger positive and negative controls of LR growth, and in particular the systemic responses of the SpWW half, also remain to be uncovered. We speculate that these processes do not involve a major transcriptional step and thereby may have escaped our analyses. (Fig. S11). For this, we are currently conducting fine analyses of xylem and phloem sap composition in root halves of plants under split WD. Finally, our work opens up important prospects for breeding crops adapted to PRD and thus making the best use of micro irrigation.

## Materials and Methods

### Plant materials and growth conditions

Seeds of maize B73-UH007 hybrid (B73h) (51) were germinated as described by Protto et al. 2024 (52), and further grown in a controlled environment growth chamber at 65% relative humidity and 22°C day/20°C night, with 16 h/8 h light/dark cycles (150 μE m^-2^ s^-1^). At 5 days after sowing (DAS), seedlings were transferred to hydroponic containers filled with 24L hydroponic solution as described in Protto et al. 2024 (52). To prevent hypoxia, culture solutions were constantly bubbled with air. High molecular weight polyethylene glycol 8000 (PEG-8000) was used to reduce the water potential and induce WD. For split root experiments, the primary root of seedlings was cut off, after 2 days of growth in a standard hydroponic solution. Seedlings were further grown for 2 days in the standard hydroponic solution to permit development of the seminal roots. At 9 DAS, 2 seminal roots of similar length were kept and plants were transferred in a homemade split-root culture system, consisting of a 24 L container with a middle separating wall allowing application of different hydroponic solutions in the two halves of the container.

For local water-deficit experiments, three different conditions were set up with distinct water availability combinations, namely Well Watered (WW) where the two halves of root systems were grown in a standard hydroponic solution, Water Deficit (WD) where the two halves of root system were grown in a standard hydroponic solution supplemented with 150 g L^-1^ PEG. The third conditions consist of plants where one seminal root is exposed to a standard hydroponic solution (Split Well Watered, SpWW) whereas the other seminal root is exposed to a PEG-induced WD (Split Water Deficit, SpWD) (Fig. 1a). Methyl jasmonate (MeJA) and 1-aminocyclopropane1-carboxylic acid (ACC) were used under homogeneous or heterogeneous root application. In homogeneous root treatment, MeJA or ACC were added at the indicated concentration into the standard hydroponic solution or the PEG hydroponic solution at 9 DAS. For split-root treatments, 50 μM MeJA was added to the standard hydroponic solution either on both sides or on one side of the split-root container, also at 9 DAS. Similar to split PEG treatments, the four different treatments were named CK, MeJA, SpCK and SpMeJA respectively.

For hydroponic experiments with *Arabidopsis thaliana* Col-0 and *jar1-1* (40), seeds were sown on a hydroponic medium (52) solidified with 0.7% Agar in black Eppendorf lids and vernalized at 4 °C in the dark. After 2 days, the holder of black Eppendorf lids was put in a 2L black container filled with a hydroponic solution. These black containers were transferred into a controlled environment growth chamber at 65% relative humidity and 25°C day/20°C night, with 8 h/16 h light/dark cycles and a light intensity of 200 μE m^-2^ s^-1^. Three weeks after sowing, the primary root was pruned below the second visible LR and plants were transferred in fresh hydroponic medium. Four days later, seedlings were transplanted into a 2L split-root container and grown for 4 days under homogeneous WW or WD conditions or under a heterogenous WD, as described for maize. For subsequent in vitro culture in plates, seeds were surface sterilized for 10 min in 1 mL of 50% (v/v) ethanol containing 2% (w/v) Bayrochlor (Bayrol, Mundolsheim, France), followed by five washes with absolute ethanol and dried in a laminar flow hood. Sterilized seeds were sown with a sterile toothpick in 12 × 12 cm transparent plates on 50 mL of solid medium (see below). After 2 days vernalization at 4 °C in the dark, plates were incubated vertically in an Aralab incubator at 65% relative humidity and 21.5°C, with a 16 h/8 h light/dark regime. Culture medium contained 2mM KNO_3_, 0.5 mM CaSO_4_, 0.5 mM MgCl_2_, 1 mM KH_2_PO_4_, 2.5 mM 2-[morpholino] ethanesulfonic acid (MES) pH 5.8, 50 μM NaFeEDTA, 50 μM H_3_BO_3_, 12 μM MnCl_2_, 1 μM CuCl_2_, 1 μM ZnCl_2_ and 0.03 μM NH_4_MoO_4_. Medium was adjusted to pH 5.7 with 1 N KOH and solidified with 1% (w/v) Type E bacto Agar. All chemicals and hormones were provided by Merk (St Quentin Fallavier, France).

### RSA Analysis

For maize and Arabidopsis grown in hydroponics, root systems were excised at harvesting time and stored in 70% ethanol. Each axial root system was manually separated on a 240 × 240 mm^2^ Petri dish containing water, and scanned with an Epson Perfection V850 Pro scanner (Epson Europe BV). All the roots were imaged as soon as possible and RSA was analyzed using the WinRhizo image analysis software (WinRhizo STD 1600, Instruments Régent Inc., Canada). Root length was quantified throughout each axial root. Several parameters were measured: axial root length, total LR length, average LR length, number of LRs. Average LR length is total LR length / number of LRs; LR density is number of LRs / axial root length.

For Arabidopsis grown in vitro, plates were scanned with an Epson Perfection V850 Pro scanner (Epson Europe BV) and RSA was analyzed using the OPTIMAS image analysis software (Adept Turnkey Pty Ltd.). Root length was extensively quantified on each root system. Several parameters were measured: primary root length, total LR length, number of LRs, average length of LRs, and LR density (number of LRs / primary root length).

### RNA extraction, sequencing and quantitative Real-time PCR Analysis

For local WD experiments or hormonal (ACC, MeJA) treatments in maize, the LRs, unbranched axial root from apex to the first visible LR (Tip) and shoots of seedlings were sampled before treatment (Control) or at the indicated time after treatment, immediately frozen in liquid nitrogen and stored at -80°C. For local WD experiments in Arabidopsis, seedlings were sampled before treatment and at 1h, 2h and 4h after treatments, separating shoots and roots and samples were immediately frozen in liquid nitrogen.

Total RNA was extracted using a Direct-zol RNA Miniprep Plus kit according to the manufacturer’s instructions (Zymo Research) with a modification for on-column DNase I digestion with addition of 5 units RQ1 RNase-Free DNase (Promega) and 1μl of a RQ1 10×buffer before downstream sequencing or RT-qPCR.

For reverse transcription quantitative real time polymerase chain reaction (RT-qPCR), 1 μg of total RNA was used as a template for first strand cDNA synthesis, which was performed using Moloney Murine Leukemia Virus Reverse Transcriptase, RNase H Minus, Point Mutant (Promega) and Oligo(dT)15 Primer (Eurofins) in a final volume of 25 μL, according to the manufacturer’s instructions. First strand cDNA was diluted 5 times, and 4μl of cDNA were used as a template for qPCR. This was performed in 384-well plates with a LightCycler 480 Real-Time PCR System (Hoffmann-La Roche AG). An intercalating dye-based qPCR master mix TB Green Premix Ex Taq (TAKARA) was used to monitor cDNA amplification. The PCR cycle program was 1 cycle of pre-denaturation for 3 min at 95°C for DNA polymerase activation, followed by 40 cycles of 5 s at 95°C and 60 s at 60°C. The 2^-ΔΔCt^ method was used to analyze the relative expression of candidate genes. The reference gene β*-ACTIN 2* (*Zm00001d012277*) and *AtUBC21* (*At5g25760*) were used to normalize gene expression in maize and *Arabidopsis*, respectively. Primer sequences are shown in Supplementary Table S5.

For sequencing, cDNA libraries were prepared and sequenced at Novogene UK Sequencing Center (Cambridge, UK) using the Illumina NovaSeq 6000 Sequencing System; 150-bp paired-end sequences with a minimum of 50 million reads were acquired for each sample. The RNA-seq data are available from NCBI Sequence Read Archive database under BioProject PRJNA1377694.

### RNA-seq data analysis

Clean RNA-seq reads were mapped to *Zea mays* reference genome B73_RefGen_v4.49 (https://nov2020-plants.ensembl.org/Zea_mays) using STAR-2.7.10a (53). HTSeq was used to count the numbers of reads mapped to each gene. Package “edgeR” was used to normalize raw counts by both library size and composition and calculate counts per million (CPM). R packages “factoextra” was used to conduct principal component analysis (PCA) using normalized CPM for all samples, LR and Tip, respectively. The package “moanin”, which models time-course data using functions and works with them, was used to detect and estimate overall differential expression. Splines were fitted to each gene using the model ∼ Treatment + Treatment: splines::ns(Time, df=4), and significant genes were selected based on FDR ≤0.01 and |log2FoldChange| ≥ 1. Splines clustering based on “kmeans” was performed and the result was visualized by function “plot_splines_data” in package “moanin” (54). “DESeq2” (55) was then used to detect differentially expressed genes (DEGs) under individual time point in more details, setting the threshold at FDR ≤ 0.01 and |log2FoldChange|≥ 1. For every DEG, CPM was used to plot heatmap using the “pheatmap” package with a z-score normalized among all LR/Tip samples. Enriched Gene Ontology terms of each group were analyzed using “gProfiler” based on Ensembl Genomes 49 (56). Gene regulatory networks were built using “DIANE” (57) based on a transcription factor list combining the maize transcription factors of “planttfdb” (http://planttfdb.gao-lab.org/index.php?sp=Zma) and the transcription factors filtered with “BioMart” under the EnsemblPlants database “Ensembl Plants Genes 49 -*Zea mays* genes (B73 RefGen_v4)” (Supplemental Table S3).

### Statistical Analyses

All statistical analyses were conducted using R v.4.2.0 (R Core Team, 2021), and figures were produced using the “ggplot2” package (58). Significance testing was performed using either two-tailed Student’s *t*-test or one-way ANOVA followed by FDR Post-Hoc Tests for multiple tests. A threshold of *p*-value < 0.05 was used in all cases, as indicated by asterisks or different letters, in Student’s *t*-tests or ANOVA, respectively.

## Acknowledgments

This work was supported by the European Research Council (ERC) under the European Union’s Horizon 2020 research and innovation programme (Grant Agreement ERC-2017-ADG-788553).

---

**Figure S1.**
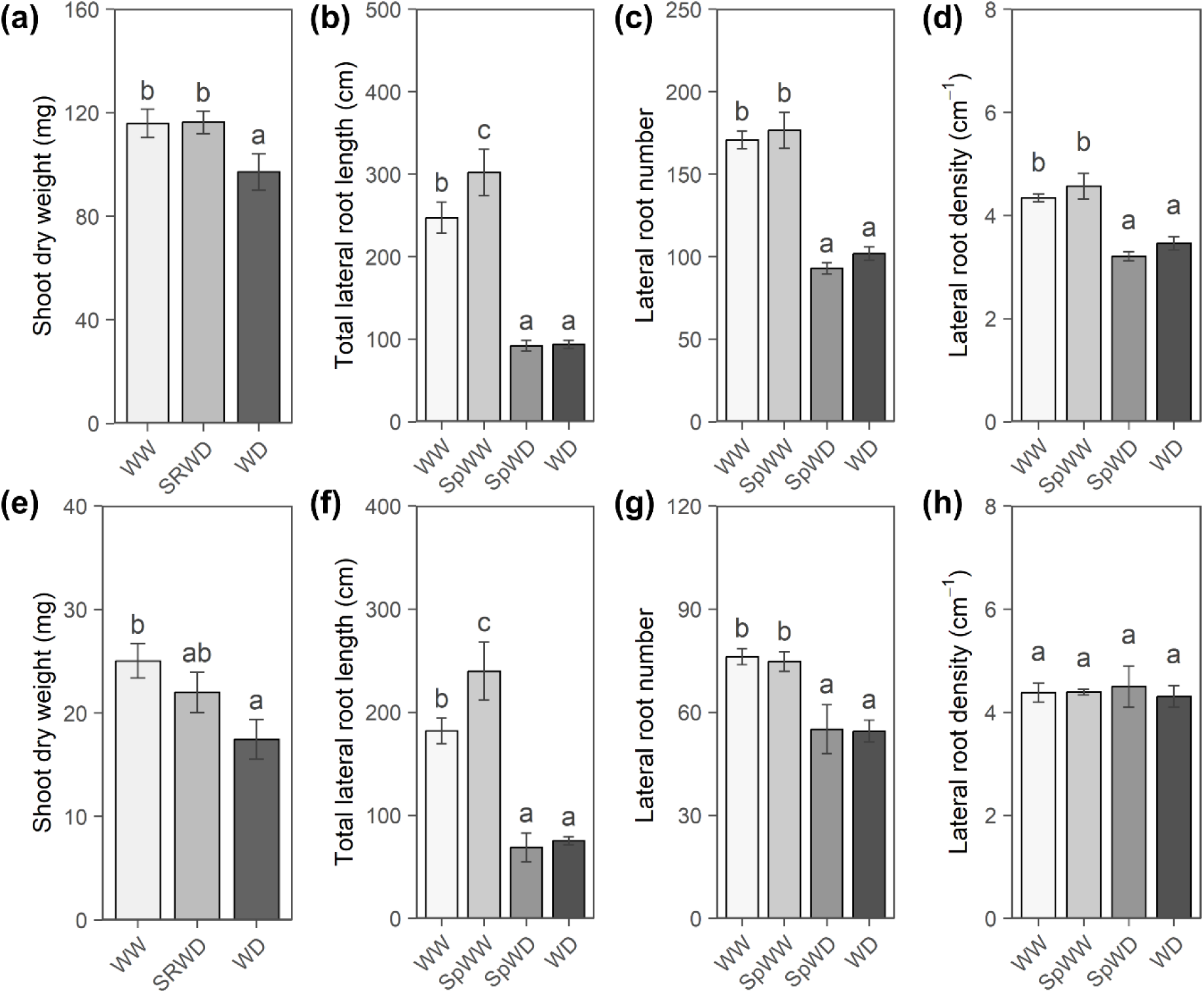
Growth responses of maize and *Arabidopsis* seedlings to a PEG-induced local WD. (**a**) Shoot dry weight of maize grown for 3 days under the following growth conditions. WW: plants with whole root system grown in a standard hydroponic solution; WD: plants with whole root systems grown in the standard hydroponic solution supplemented with 150 g L^-1^ PEG; SRWD: plants grown in the split root system with one half of the roots in the standard solution (SpWW) and the other half in the standard hydroponic solution containing 150 g L^-1^ PEG (SpWD); (**b-d**) LR growth of WW, SRWD and WD maize plants after 3 days of treatment; (**e**) Shoot dry weight of WW, SRWD and WD Arabidopsis plants after 4 days of treatment; (**f-h**) LRs growth of WW, SRWD and WD Arabidopsis plants after 4 days of treatment. The bar plots show cumulated data from 3 independent experiments (n=16-18 plants). All data represent mean ± SE. For each plot, different letters indicate statistically different values (P < 0.05, one-way ANOVA followed by FDR test).

**Figure S2.**
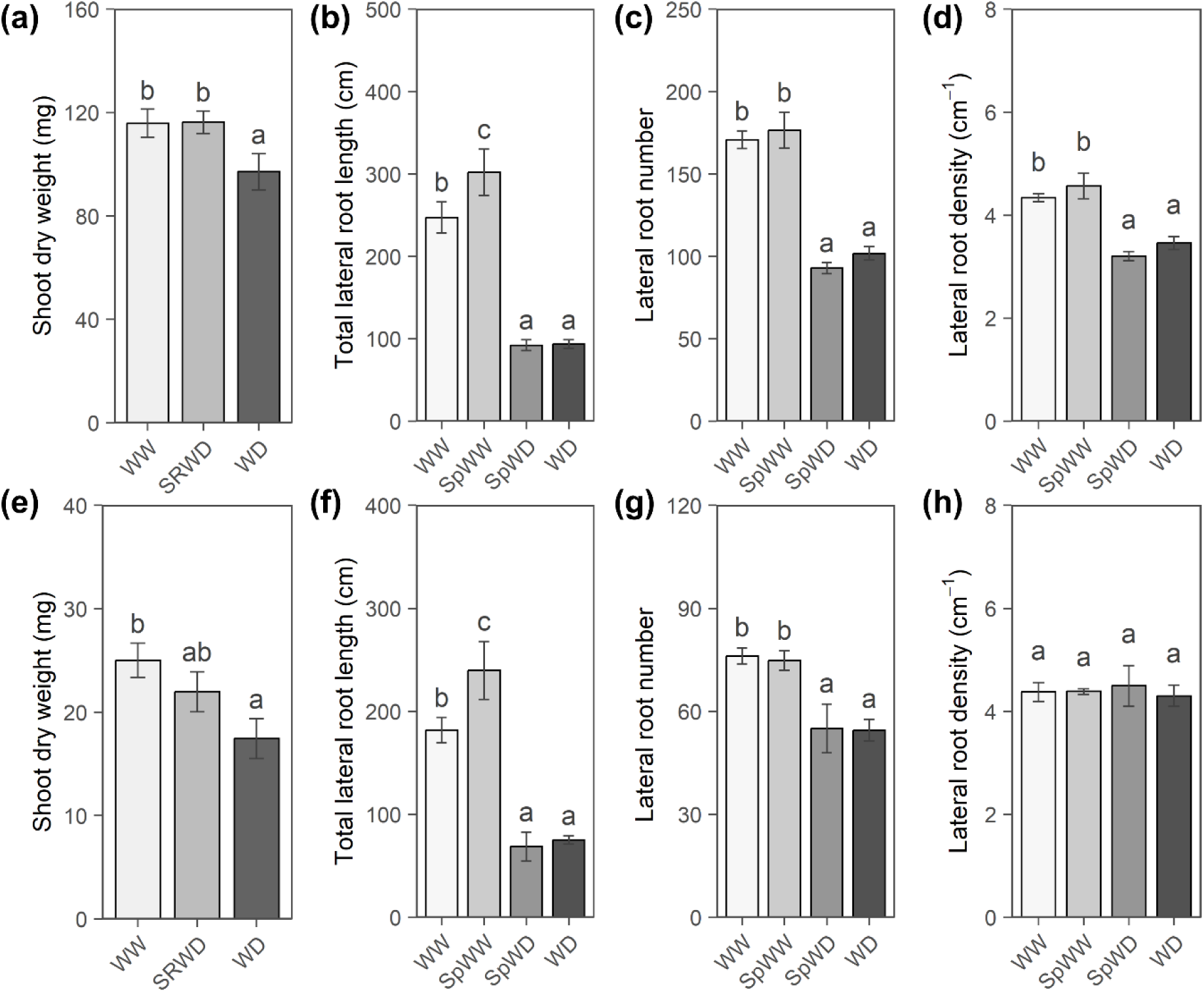
Systemic effects of a 5-day-long local WD on LR elongation in maize. (**a, b**) Shoot and root dry weight of WW, SRWD, and WD maize plants after 5 days of treatment. Experimental conditions and figure conventions for local WD are as in Figure S1. Cumulated data from 2 independent experiments (WW: n = 29, SpWW: n = 20, SpWD: n = 13, WD: n = 22). (**c**) Axial root length of WW, SRWD, and WD maize plants normalized to WW plants. (**d, e**) Relative length and number of LRs in WW, SRWD, and WD maize plants as above. For figures **c-e**, cumulated data from 4 independent experiments (WW: n = 57, SpWW: n = 50, SpWD: n = 32, WD: n =40). Error bars indicate SE. For each root type, different letters indicate statistically different values (ANOVA followed by FDR test; *P* < 0.05). Asterisk indicates significant difference after a Student’s *t* test (*P* < 0.05).

**Figure S3.**
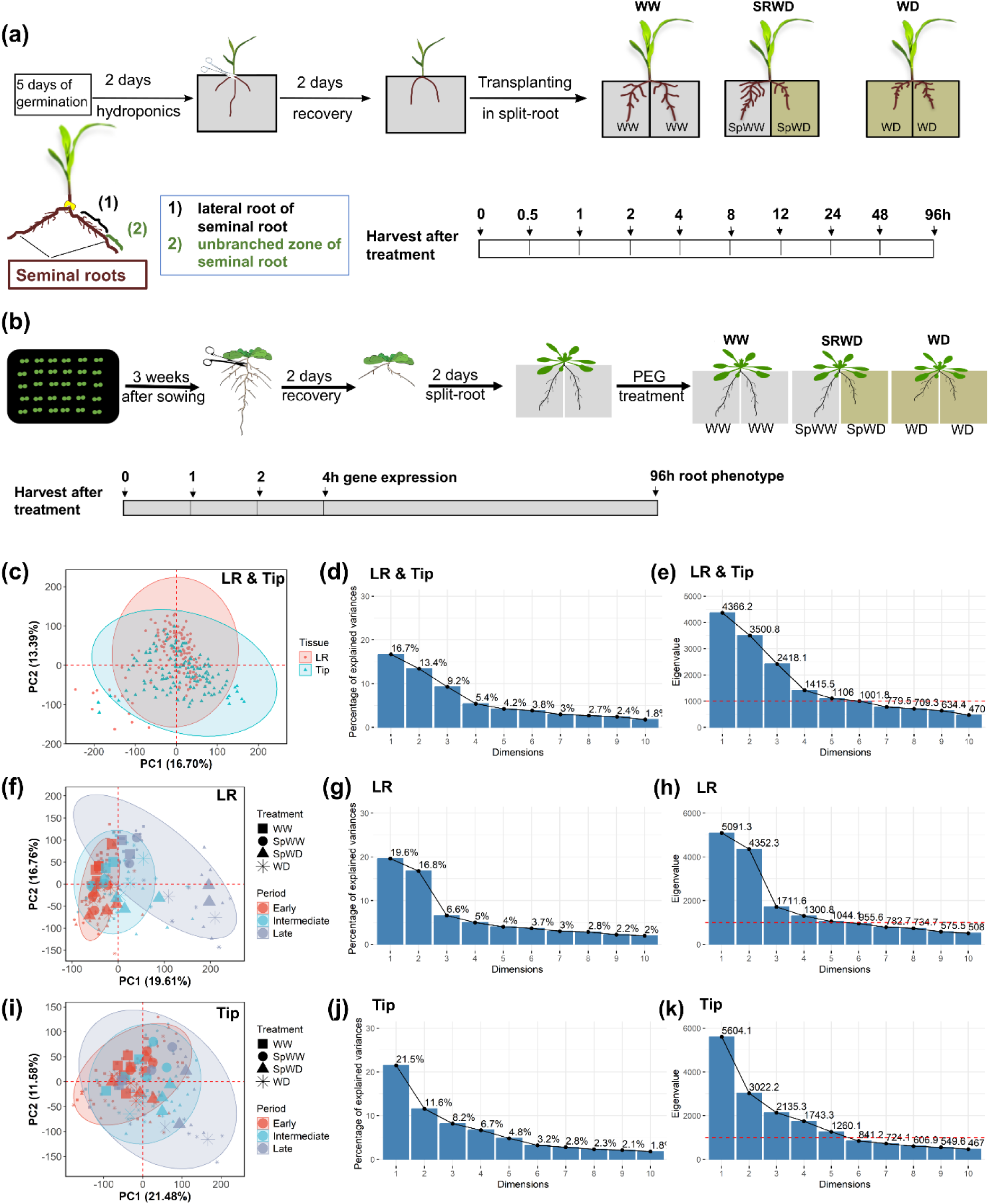
Global transcriptional response of LR and Tip tissues to a local WD. (**a**) Experimental design of transcriptome time-series to explore the effects of a local WD on gene expression in maize roots. Plants were harvested over 10 time points in two fractions LR and Tip. Four independent biological replicates were collected to perform RNA-seq. (**b**) Experimental procedure to explore the effect of a local WD on gene expression in Arabidopsis roots (**c**) PCA results of temporal gene expression in all samples of maize (**d**) Scree plot of proportion of variance explained by the top 10 components of PCA; (**e**) Scree plot of the eigenvalue determined in the top 10 components of PCA; (**f**) PCA results of temporal gene expression in LR samples of maize. (**g, h**) Scree plots of proportion of variance and of the eigenvalue as defined in (**d**) and (**e**) but for LR samples; (**i**) PCA results of temporal gene expression in Tip samples (**j, k**) Scree plot of proportion of variance and of the eigenvalue for Tip samples, as defined in (**d**) and (**e**). Large symbols in (**f**) and (**I**) represent the mean of the four samples.

**Figure S4.**
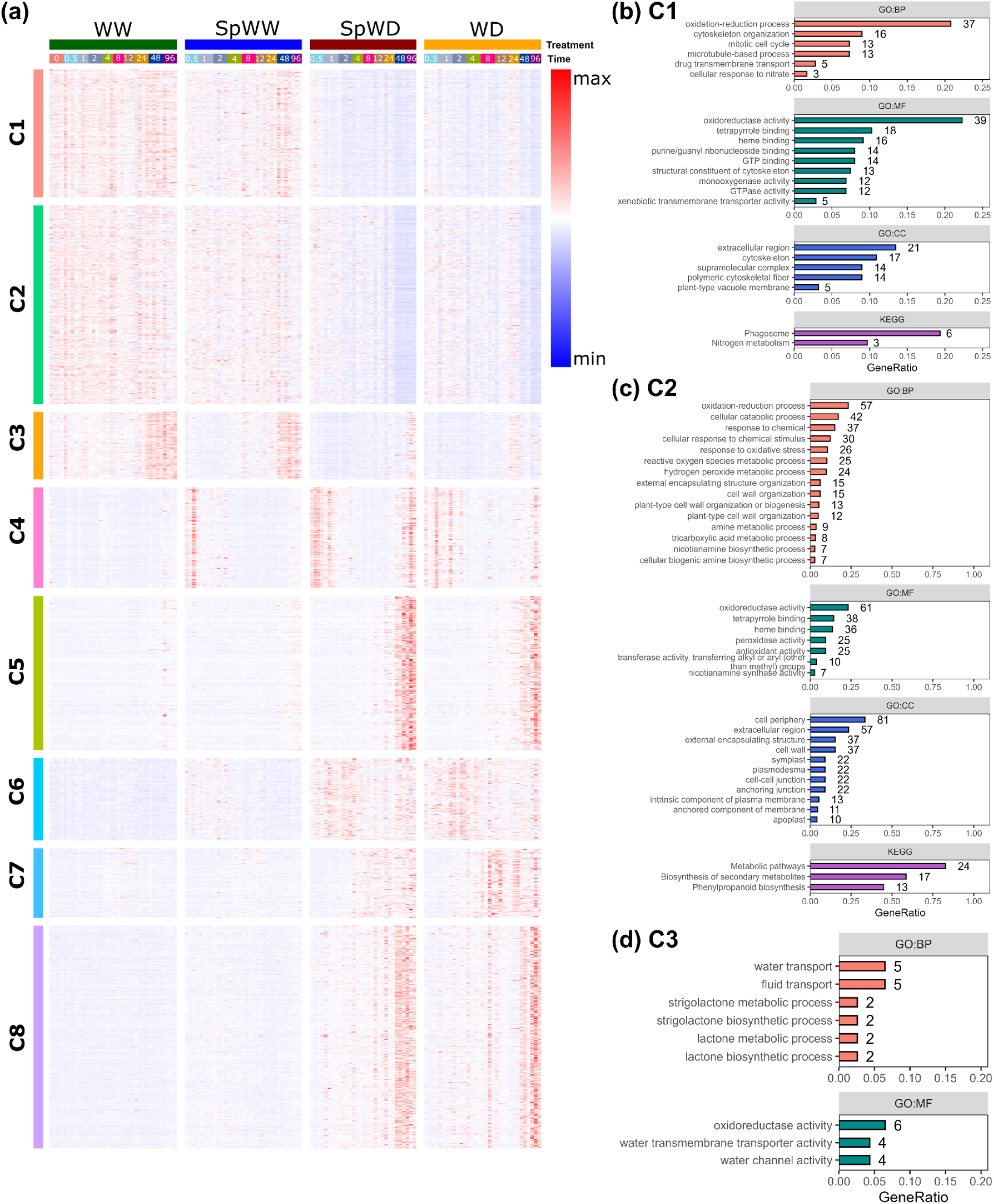
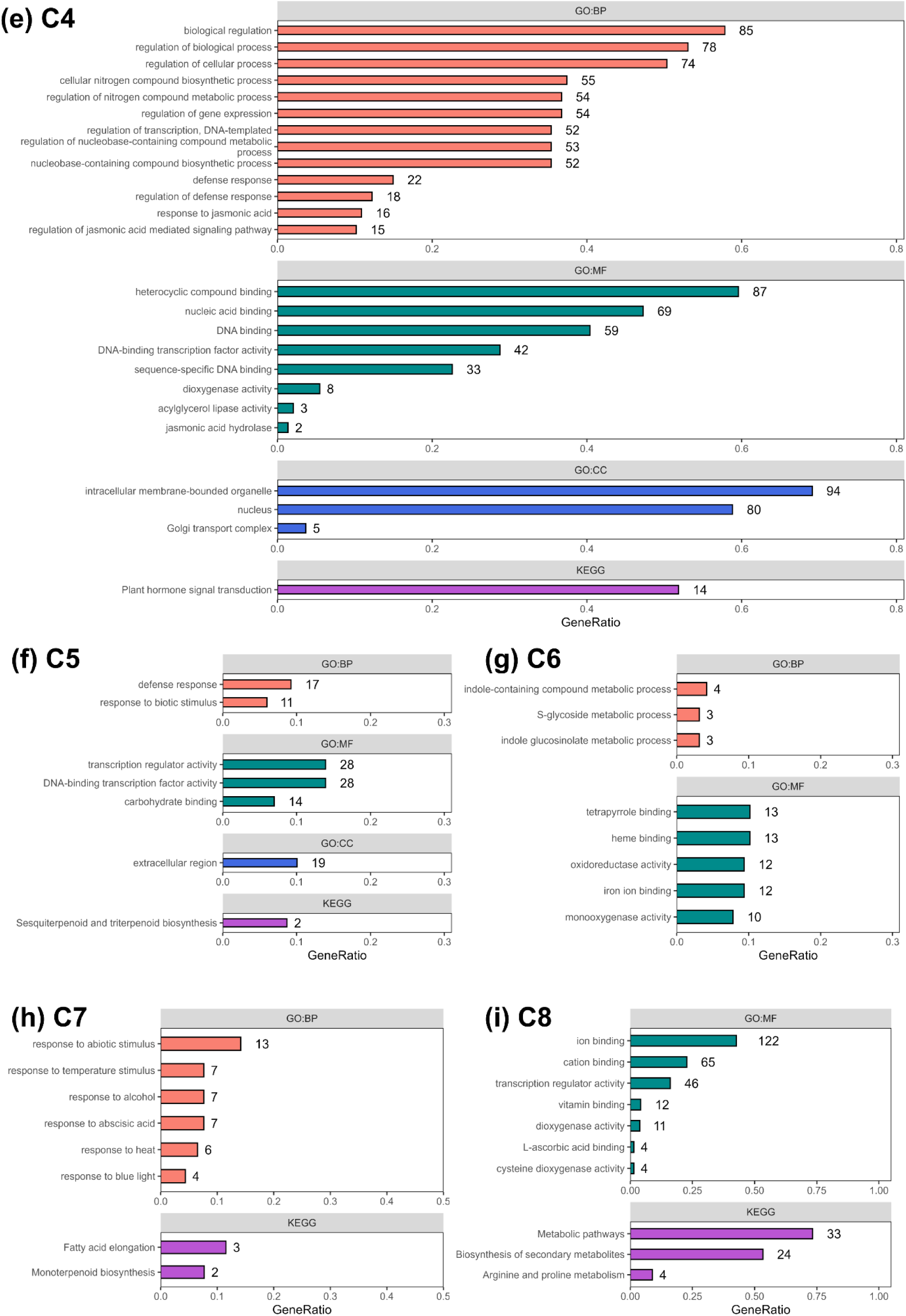
Effects of a local WD on gene expression in LR samples of maize seedlings. (**a**) The expression heatmap and k-means clustering of DEGs identified in the LR response to a local WD reveals eight major gene clusters (C1-C8). The columns of the figure represent gene expression under WW, SpWW, SpWD, and WD conditions, respectively. Each column is subdivided into time points, which themselves contain the individual replicates. Note a strong and transient induction of genes expression at early time points in Cluster 4 in all but WW conditions. (**b-i**) Gene Ontology (GO) and KEGG pathway representation analysis of DEGs in clusters C1-C8. The term names were ranked from top to bottom according to the gene ratio (BP: biological process; MF: molecular function; CC: cell component).

**Figure S5.**
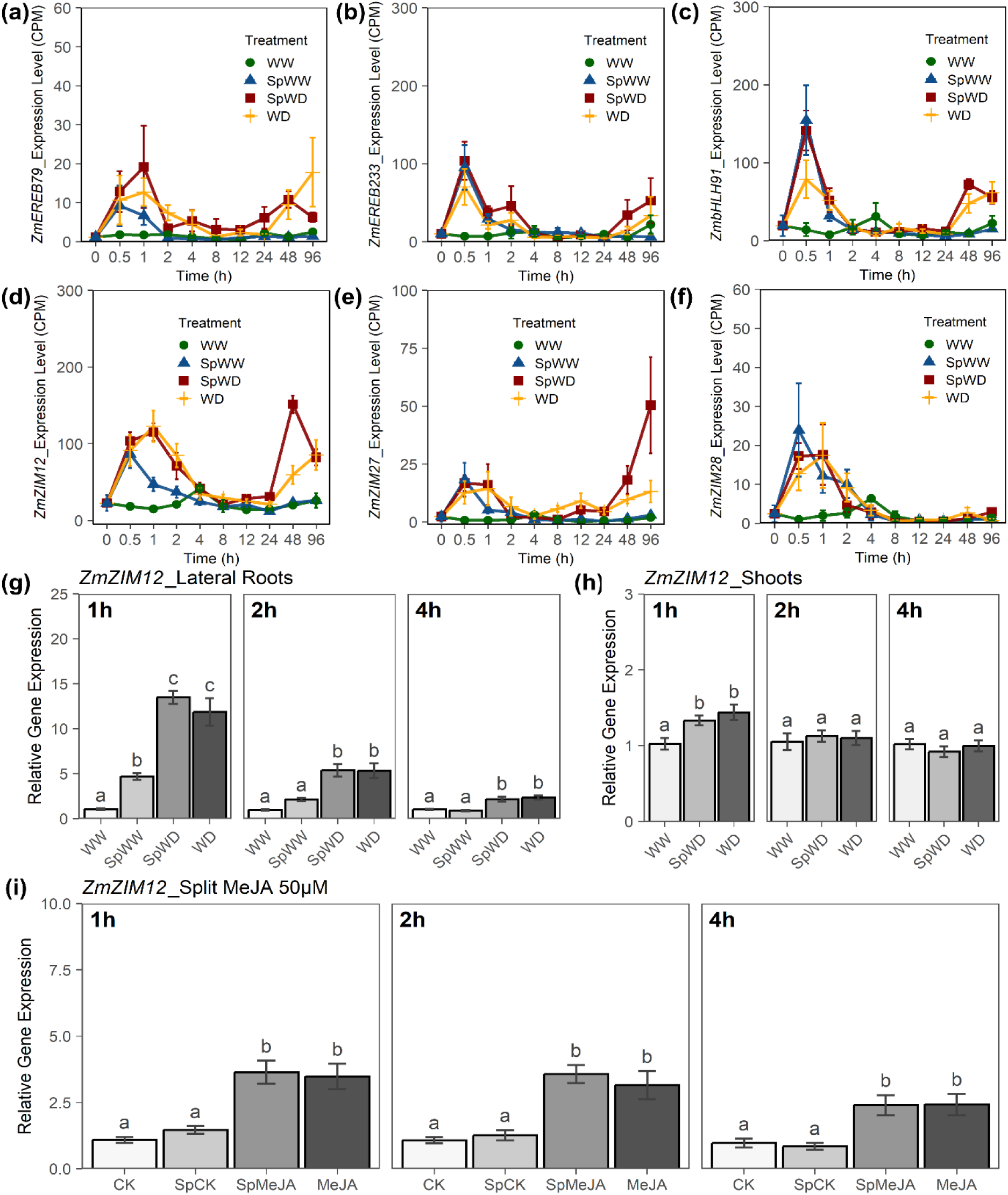
Expression pattern of JA responsive genes in maize. (**a-f**) Expression pattern from RNA seq experiments, of JA responsive genes *ZmEREB79*, *ZmEREB233*, *ZmbHLH91*, *ZmZIM12*, *ZmZIM27* and *ZmZIM28* in LRs of plants under control conditions (WW), or after a homogeneous or a local WD. **(g-h**) *ZmZIM12* expression in LRs and shoot in response to a local WD; (**i**) *ZmZIM12* expression in LRs after a short term homogeneous or local MeJA (50uM) application. Experimental conditions and figure conventions for local WD are as above. For split-root MeJA treatment, CK and MeJA correspond to plants with their two root halves in a standard hydroponic solution or in hydroponic solution supplemented by 50 µM MeJA, respectively. For heterogeneous MeJA treatments, plants were grown with one root half in the standard solution (SpCK) and the other half in a standard hydroponic solution supplemented by 50 µM MeJA (SpMeJA). All data represent mean ± SE of 4 biological replicates with 2 technical repeats; different letters indicate statistically different values (P < 0.05, one-way ANOVA followed by FDR test)

**Figure S6.**
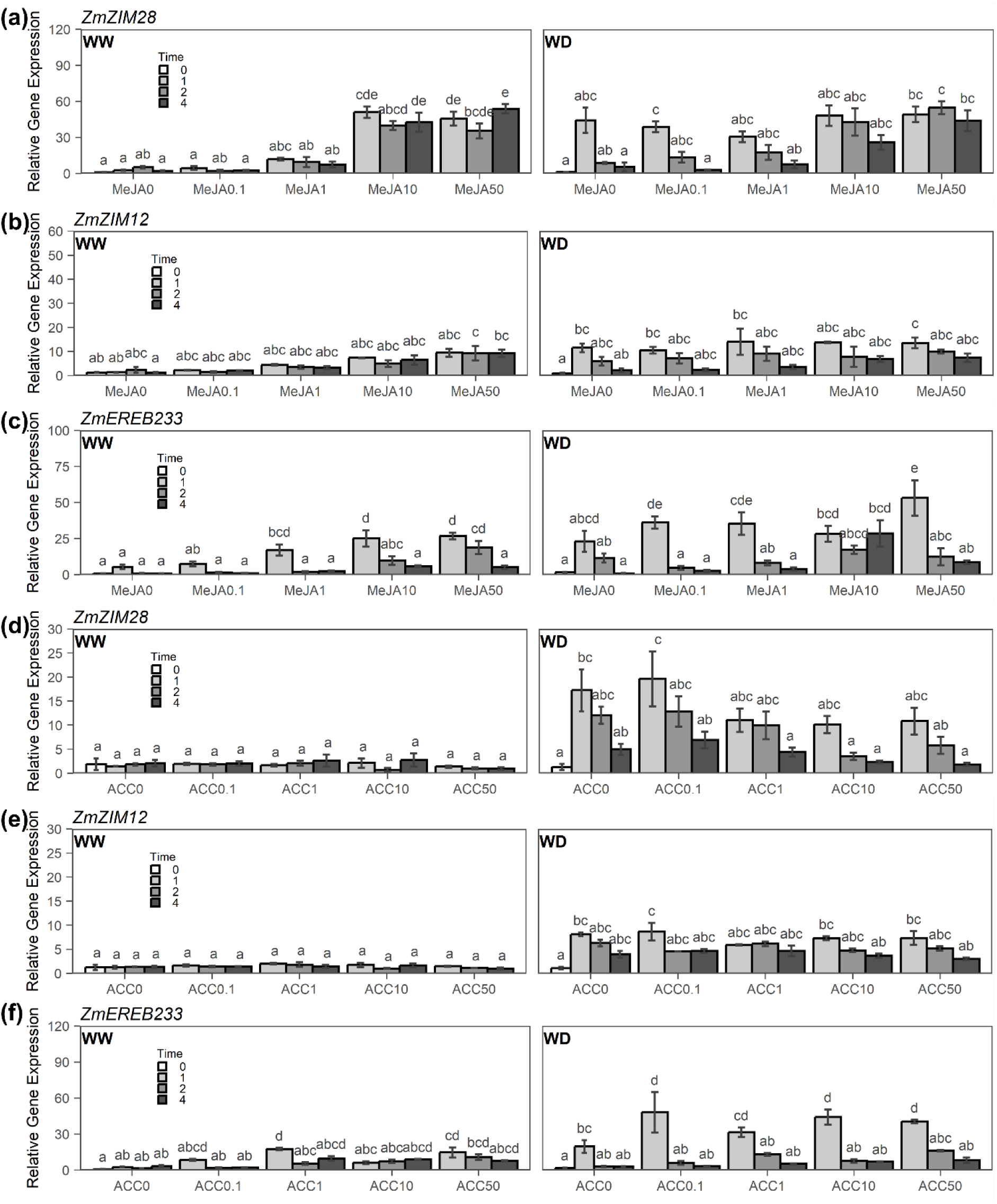
Expression of JA and ET responsive genes in LRs of maize. (**a-c**) *ZmZIM28*, *ZmZIM12* and *ZmEREB233* expression pattern following short term exposure to increasing MeJA concentrations under WW or WD conditions (**e-f**) *ZmZIM28*, *ZmZIM12* and *ZmEREB233* expression pattern following short term exposure to increasing ACC concentrations under WW or WD conditions The MeJA and ACC were both applied at 0 μM, 0.1 μM, 1 μM, 10 μM or 50 μM. Data represent mean ± SE of 3-4 biological replicates with 2 technical replicates. For each time point after treatment, different letters indicate significant differences (P < 0.05, one-way ANOVA, FDR test).

**Figure S7.**
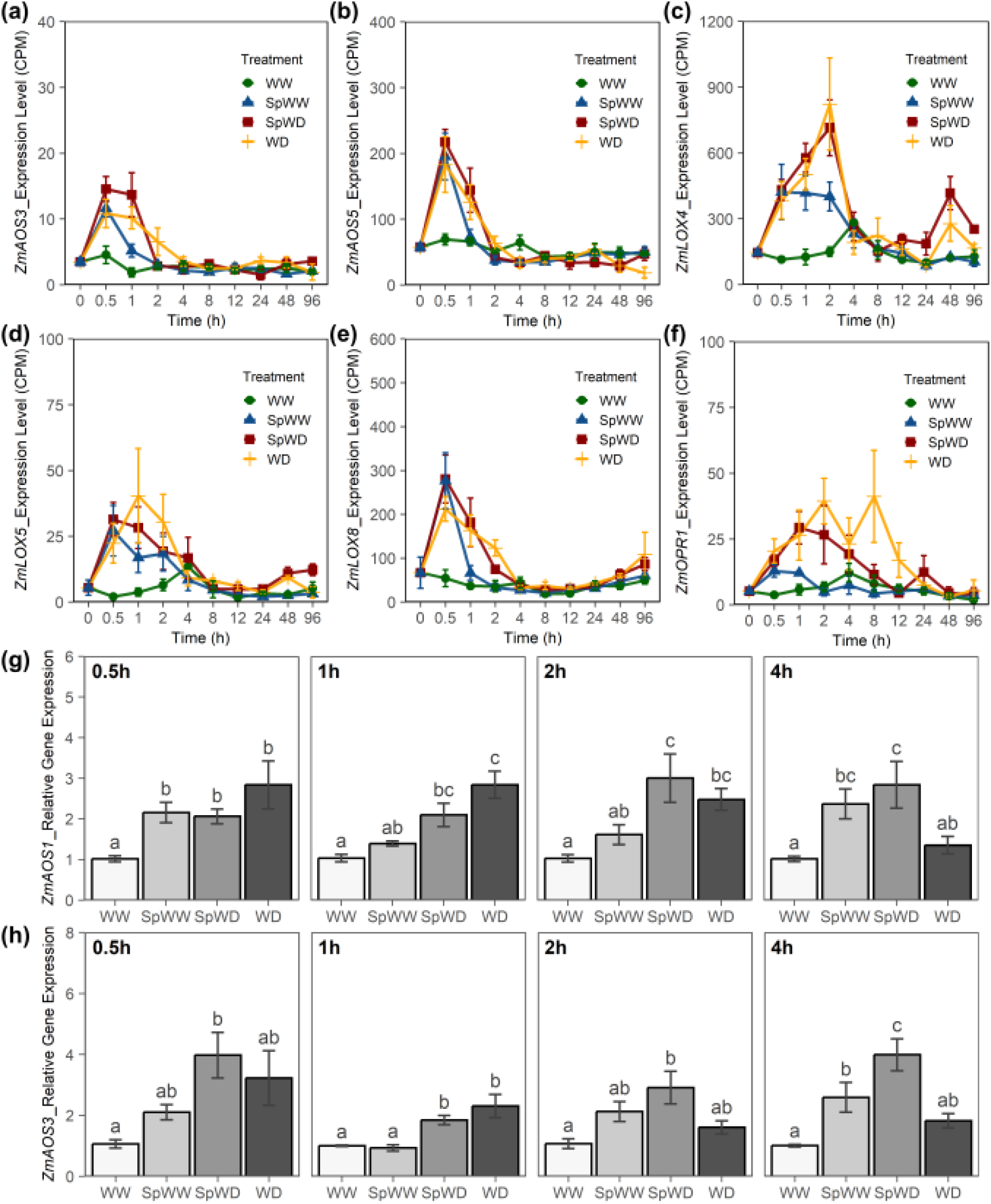
Expression pattern of JA synthesis genes in maize LRs. (a-f) Kinetic expression profile of JA biosynthesis genes, *ZmAOS3*, *ZmAOS5*, *ZmLOX4*, *ZmLOX5*, *ZmLOX8* and *ZmOPR1*, in the LR of root halves grown under control conditions (WW), or after a homogeneous or local WD; (g-h) *ZmAOS1* and *ZmAOS3* expression under WW, WD and SRWD in LRs; Experimental conditions and figure conventions for local WD are as above. The bar plots show cumulated data from 3-4 independent experiments (n=15-30) with 2 technical replicates. All data represent mean ± SE, different letters indicate significant differences (P < 0.05, one-way ANOVA, FDR test).

**Figure S8.**
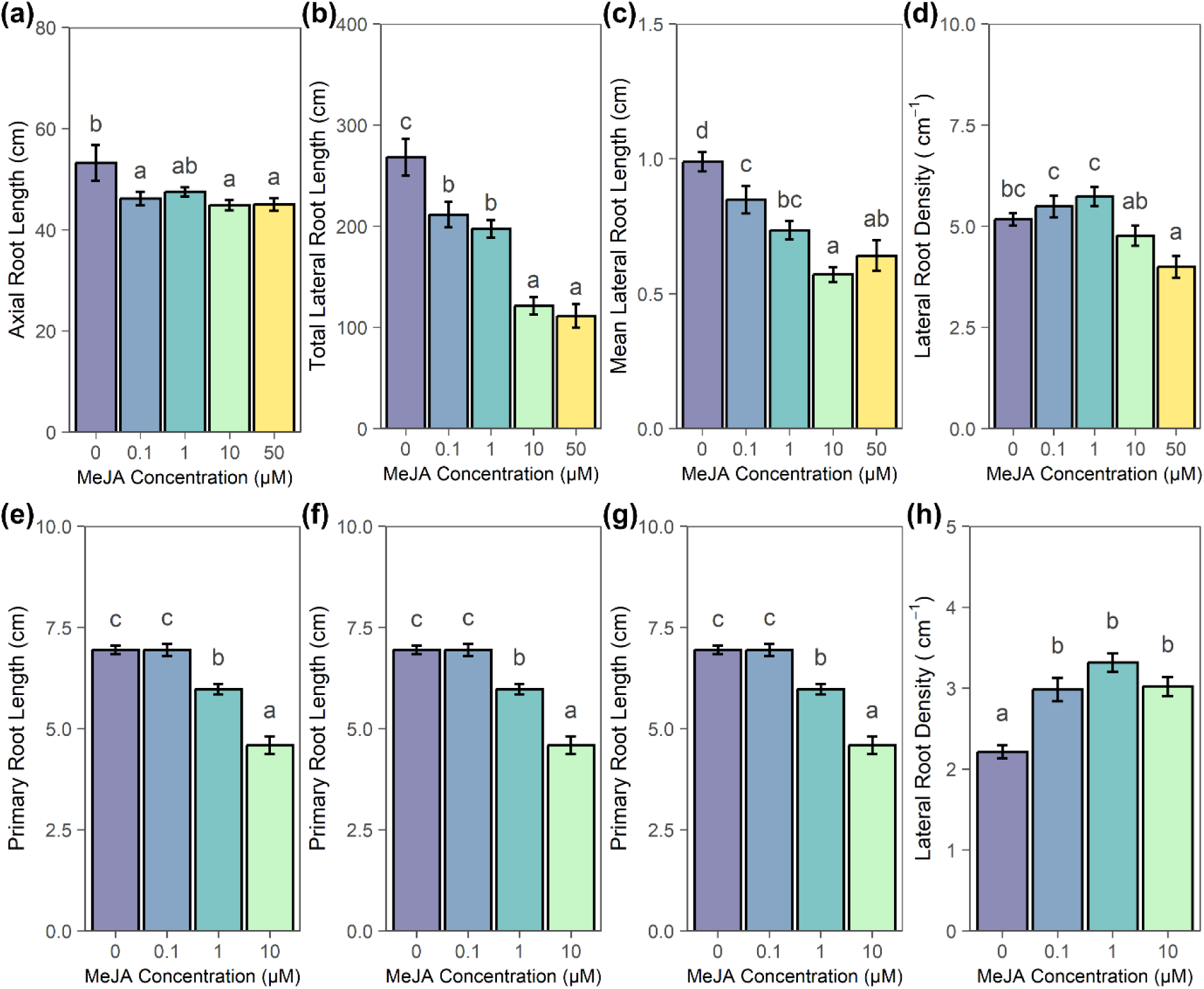
Dose-dependent effects of MeJA on maize and Arabidopsis root growth. (**a**) Axial root length of maize seedlings grown for 3 days in a standard hydroponic solution complemented with indicated MeJA concentrations; (**b-d**) LRs development parameters of maize seedlings grown as in (**a**). (**e**) Primary root length of Arabidopsis plants grown for 5 days in vertical agar plates with indicated MeJA concentrations; (**f-h**) LRs growth parameters of Arabidopsis plants grown as in (**e**). The bar plots show cumulated data from 3 independent experiments (n=12-18). All data represent mean ± SE, different letters indicate significant differences (P < 0.05, one-way ANOVA, FDR test).

**Figure S9.**
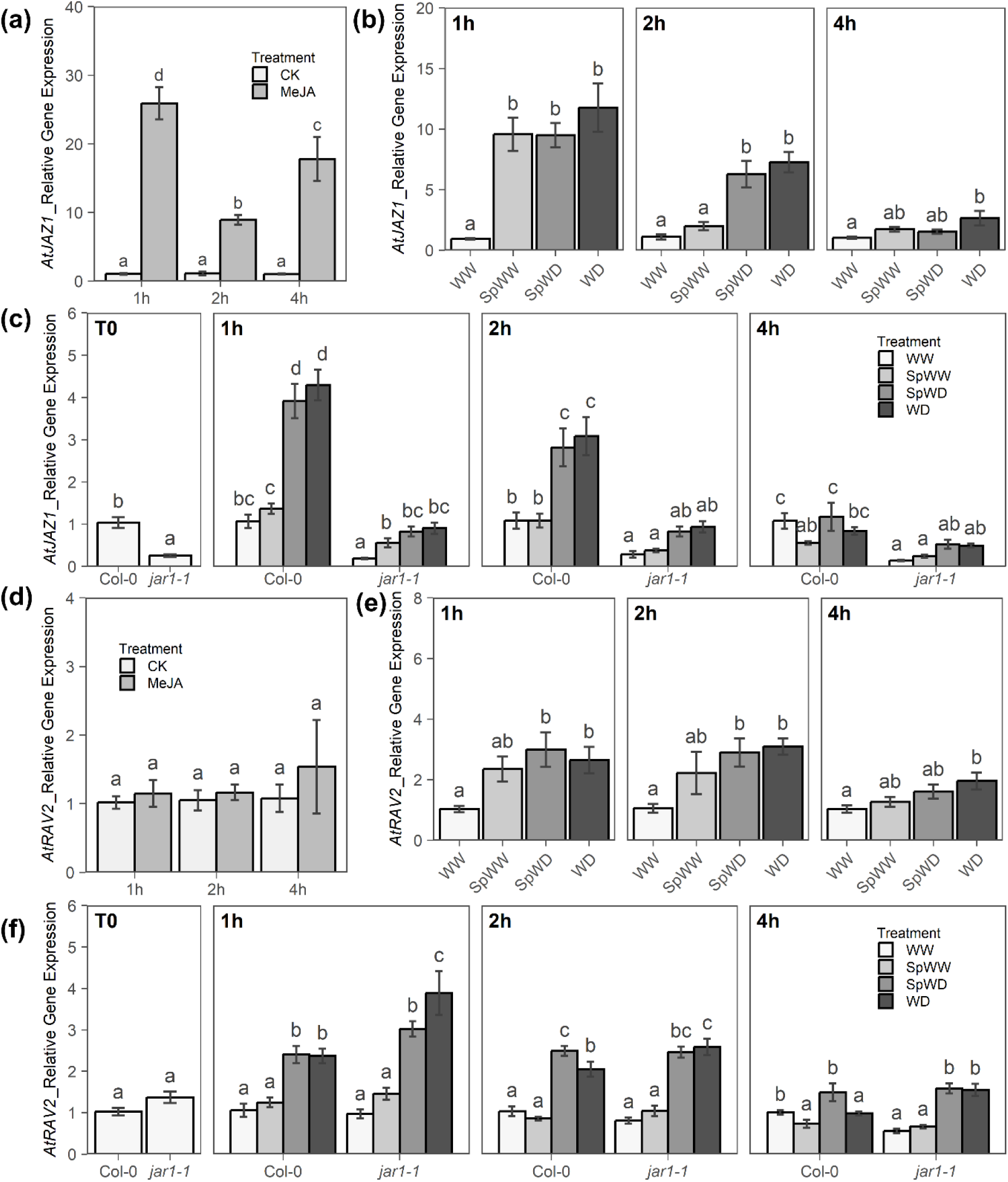
Expression pattern of JA responsive genes in *Arabidopsi*s. (**a**) Expression pattern of *AtJAZ1* in Col-0 roots in response to methyl jasmonate (50 µM -MeJA50) treatment; (**b**) *AtJAZ1* expression in Col-0 roots and at the indicated times after transfer under control (WW), or after a homogeneous or local WD; (**c**) *AtJAZ1* expression in Col-0 and *jar1-1* roots at the indicated times after transfer under control (WW), or after a homogeneous or local WD; (**d**) AtRAV2 in Col-0 roots expression in response to methyl jasmonate (50 µM -MeJA50) treatment; (**e**) AtRAV2 expression in Col-0 roots and at the indicated times after transfer under control (WW), or after a homogeneous or local WD; (**f**) AtRAV2 expression in Col-0 and *jar1-1* roots at the indicated times after transfer under control (WW), or after a homogeneous or local WD. The bar plots show cumulated data from 3-4 independent experiments (n=18-30). All data represent mean ± SE, different letters indicate significant differences (P < 0.05, one-way ANOVA, FDR test).

**Figure S10.**
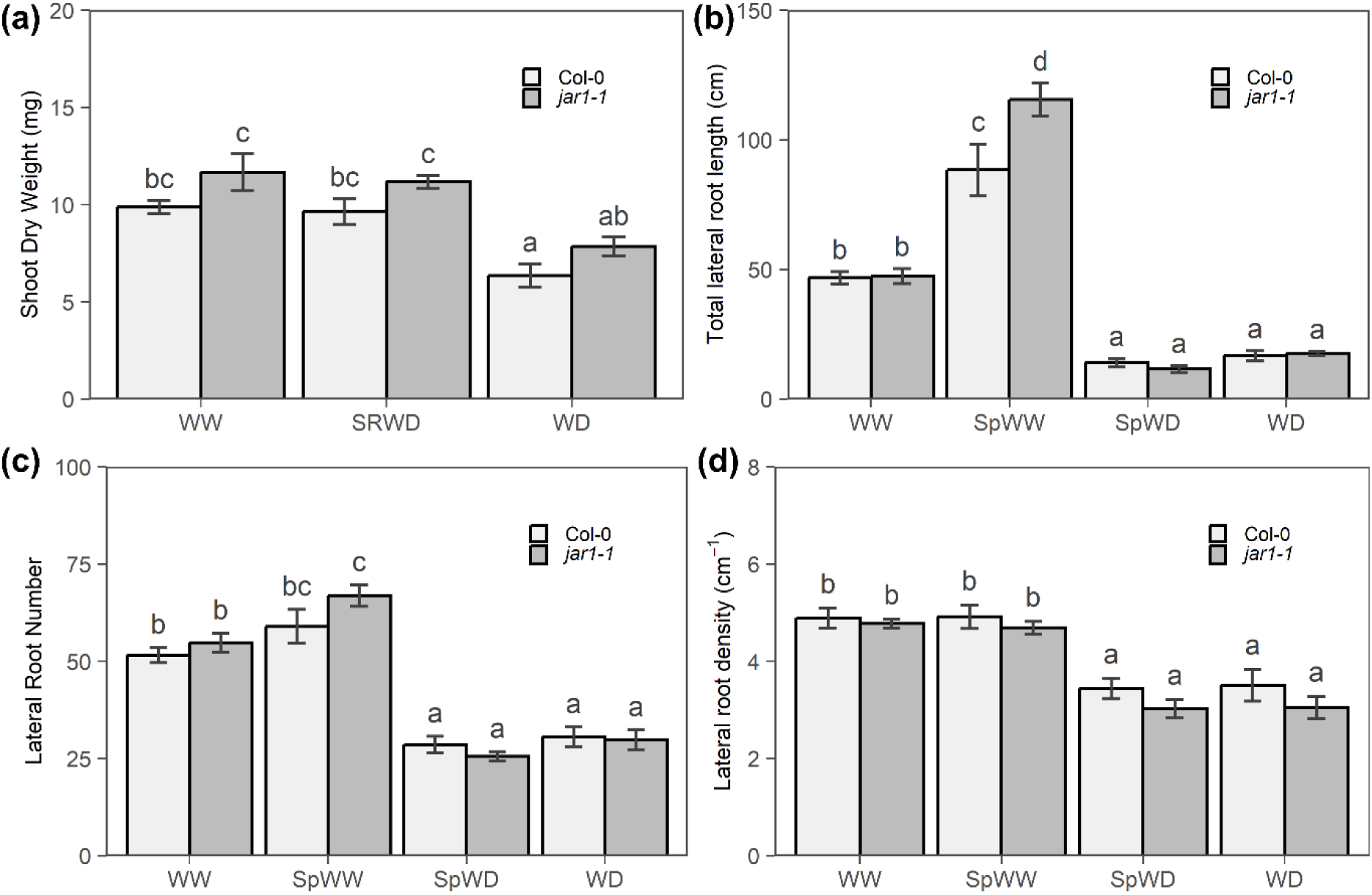
Growth responses Arabidopsis Col-0 and jar1-1 seedlings to a local WD. (**a**) Shoot dry weight (**b-d**) Total LR length, LR number and LR density after 4 days of growth. Experimental conditions and figure conventions for local WD are as above. The bar plots show cumulated data from 2 independent experiments (n=12-20). All data represent mean ± SE. For each shoot or root trait, different letters indicate statistically different values (P < 0.05, one-way ANOVA followed by FDR test).

**Figure S11.**
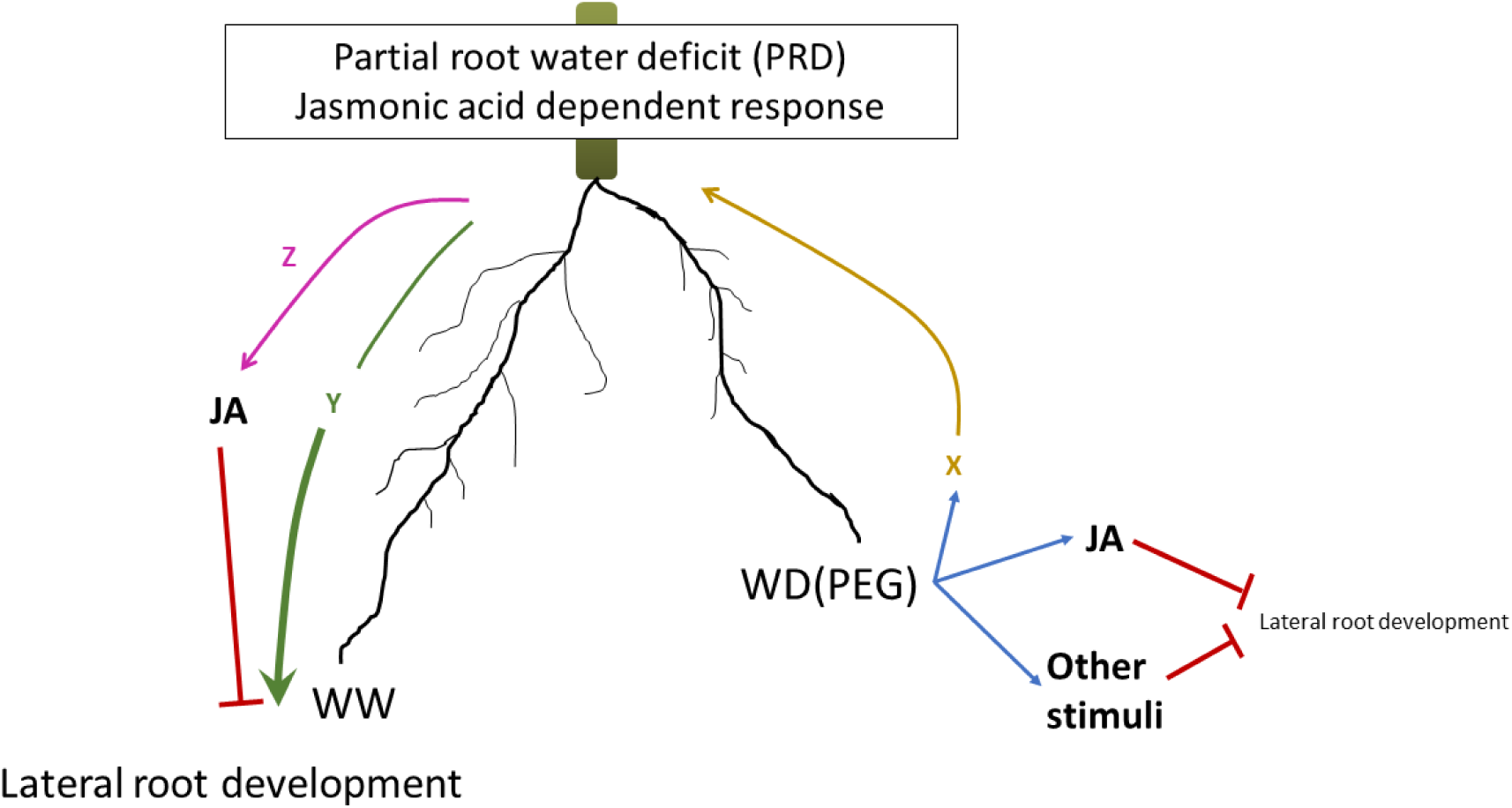
Working model of JA-dependent responses to partial-root water deficit (PRD). The figure shows how a local WD (right root half) induces the biosynthesis of JA and/or other signaling molecules which lead to inhibition of lateral root development in the WD or SpWD root halves. A signal X is also produced in SpWD root and exported towards the shoot. This signal either relay signals (Y, Z) induce JA biosynthesis or directly enhance LR development. The figure shows that locally produced JA in turns inhibits LR development, thereby counteracting the effects of Y. Note that X, Y, Z could be similar or different, and remain to be identified.

**Table S1.** Gene name and description for each DEG Cluster.

**Table S2.** Gene Ontology over-representive analysis result for each DEG Cluster.

**Table S3.** Regulators of maize used to build gene regulatory network in this study.

**Table S4.** Gene regulatory network for each DEG Cluster.

**Table S5:** RT-qPCR primers sequence.

